# ZNF410 uniquely activates the NuRD component CHD4 to silence fetal hemoglobin expression

**DOI:** 10.1101/2020.08.31.274324

**Authors:** Xianjiang Lan, Ren Ren, Ruopeng Feng, Lana C. Ly, Yemin Lan, Zhe Zhang, Nicholas Aboreden, Kunhua Qin, John R. Horton, Jeremy D. Grevet, Thiyagaraj Mayuranathan, Osheiza Abdulmalik, Cheryl A. Keller, Belinda Giardine, Ross C. Hardison, Merlin Crossley, Mitchell J Weiss, Xiaodong Cheng, Junwei Shi, Gerd A. Blobel

## Abstract

Metazoan transcription factors typically regulate large numbers of genes. Here we identify via a CRISPR-Cas9 genetic screen ZNF410, a pentadactyl DNA binding protein that in human erythroid cells directly and measurably activates only one gene, the NuRD component CHD4. Specificity is conveyed by two highly evolutionarily conserved clusters of ZNF410 binding sites near the CHD4 gene with no counterparts elsewhere in the genome. Loss of ZNF410 in adult-type human erythroid cell culture systems and xenotransplant settings diminishes CHD4 levels and derepresses the fetal hemoglobin genes. While previously known to be silenced by CHD4, the fetal globin genes are exposed here as among the most sensitive to reduced CHD4 levels. In vitro DNA binding assays and crystallographic studies reveal the ZNF410-DNA binding mode. ZNF410 is a remarkably selective transcriptional activator in erythroid cells whose perturbation might offer new therapeutic opportunities in the treatment of hemoglobinopathies.

**Highlights:**

- A CRISPR screen implicates ZNF410 in fetal globin gene repression
- The CHD4 gene is the singular direct ZNF410 target in erythroid cells
- The fetal globin genes are exquisitely sensitive to CHD4 levels
- Five C2H2 zinc fingers of ZNF410 recognize the major groove of a 14 base pair sequence

## Introduction

In bacteria, one regulatory transcription factor (TF) often controls the expression of a single gene or operon (Jacob et al., 1960). In contrast, the vast majority of mammalian TFs regulate many target genes. Spatio-temporal specificity of gene transcription is achieved by combinatorial deployment of TFs and their co-regulators. For example, the transcription factor GATA1 cooperates with the TF KLF1 and TAL1/SCL to regulate erythroid-specific gene expression (Love et al., 2014). whereas GATA1 and ETS family TFs regulate megakaryocyte-enriched genes (Wang et al., 2002). In erythroid cells, the most highly expressed genes encode the α- and β-subunits of the hemoglobin tetramer. The human β-type globin gene cluster consists of one embryonic gene (HBE, also known as ε-globin), two fetal genes (HBG1 and HBG2, also known as ^G^γ-globin and ^A^γ-globin), and two adult genes (HBB and HBD, also known as β-globin and δ-globin) genes. The ε-globin gene is transcribed in primitive erythroid cells in early development, and during mid-gestation, is silenced concomitantly with the γ-globin genes turning on. Around the time of birth a second switch occurs when β- and δ-globin transcription is activated at the expense of the γ-globin genes. Therefore, disease causing alterations in the β-globin gene such as those causing sickle cell disease (SCD) and some types of β-thalassemia become symptomatic after birth, coincident with the γ-to-β–globin switch. Reversing the switch from β-globin back to γ-globin expression in developing erythroid cells has been a major endeavor for treating these diseases (Platt et al., 1994; Wienert et al., 2018).

While lineage restricted TFs such as GATA1 and TAL1 are essential for erythroid specific transcription, two zinc-finger TFs, BCL11A and LRF (ZBTB7A), play a dominant role in the developmental control of the fetal to adult switch in globin gene transcription (Masuda et al., 2016; Menzel et al., 2007; Sankaran et al., 2008; Uda et al., 2008). Both of these factors bind at several locations along the β-globin gene cluster, including the promoter and upstream regions of the γ-globin genes to silence their expression (Liu et al., 2018; Martyn et al., 2018). Both factors interact with the CHD4/NuRD complex, and CHD4 and associated proteins are required for transcriptional repression of the γ-globin genes (HBG1/2) (Amaya et al., 2013; Masuda et al., 2016; Xu et al., 2013). Given that BCL11A contains a motif found in a variety of NuRD associated molecules that is necessary and sufficient for NuRD binding (Hong et al., 2005; Lejon et al., 2011), the most parsimonious model is that BCL11A and LRF are direct links between NuRD and the γ-globin genes. One key unanswered question that is whether the expression of NuRD proteins themselves is regulated, and whether control of NuRD expression might be part of the γ-globin regulatory circuitry.

To search for novel regulators of γ-globin expression, we screened a sgRNA library targeting the DNA-binding domains of most known human transcription factors using an optimized protein domain-focused CRISPR-Cas9 screening platform (Grevet et al., 2018; Shi et al., 2015). We found that zinc finger 410 (*ZNF410*, APA-1), a transcription factor with five tandem canonical C2H2-type zinc fingers (ZFs) is required for the maintenance of γ-globin silencing. RNA-seq, ChIP-seq and genetic perturbation led to the remarkable finding that ZNF410 regulates CHD4 as its sole direct target gene via two binding site clusters not found elsewhere in the genome. We further demonstrate that the γ-globin genes are exquisitely sensitive to CHD4 levels. DNA binding and crystallographic studies reveal the mode of ZNF410 interaction with DNA. To our knowledge, ZNF410 is the only known mammalian TF with a single regulatory target in erythroid cells.

## Results

### CRISPR-Cas9 screen identifies ZNF410 as a candidate γ-globin repressor

To identify novel regulators of HbF expression, we screened a sgRNA library containing 6 sgRNAs each targeting the DNA-binding domain of most human transcription factors (1436 total, on average 6 sgRNAs each) (Huang et al., 2020). A lentiviral vector library encoding the sgRNAs was used to transfect the human adult-type erythroid cell line HUDEP-2 that stably expresses spCas9 (HUDEP-2-Cas9; (Grevet et al., 2018)). The top 10% and bottom 10% of HbF expressing cells were purified via anti-HbF FACS, and representation of each sgRNA in the two populations assessed by deep sequencing (Figure 1A). As expected, control non-targeting sgRNAs were evenly distributed between the HbF-high and HbF-low populations. Positive control sgRNAs targeting the known γ-globin repressor genes BCL11A and LRF were enriched in the HbF-high population (Figure 1B), validating the screen. Six sgRNAs against a novel TF, ZNF410, with no known prior role in globin gene regulation were significantly enriched in the HbF-high population (Figure 1B), suggesting that ZNF410 may function as a direct or indirect repressor of γ-globin expression.

**Figure 1.**
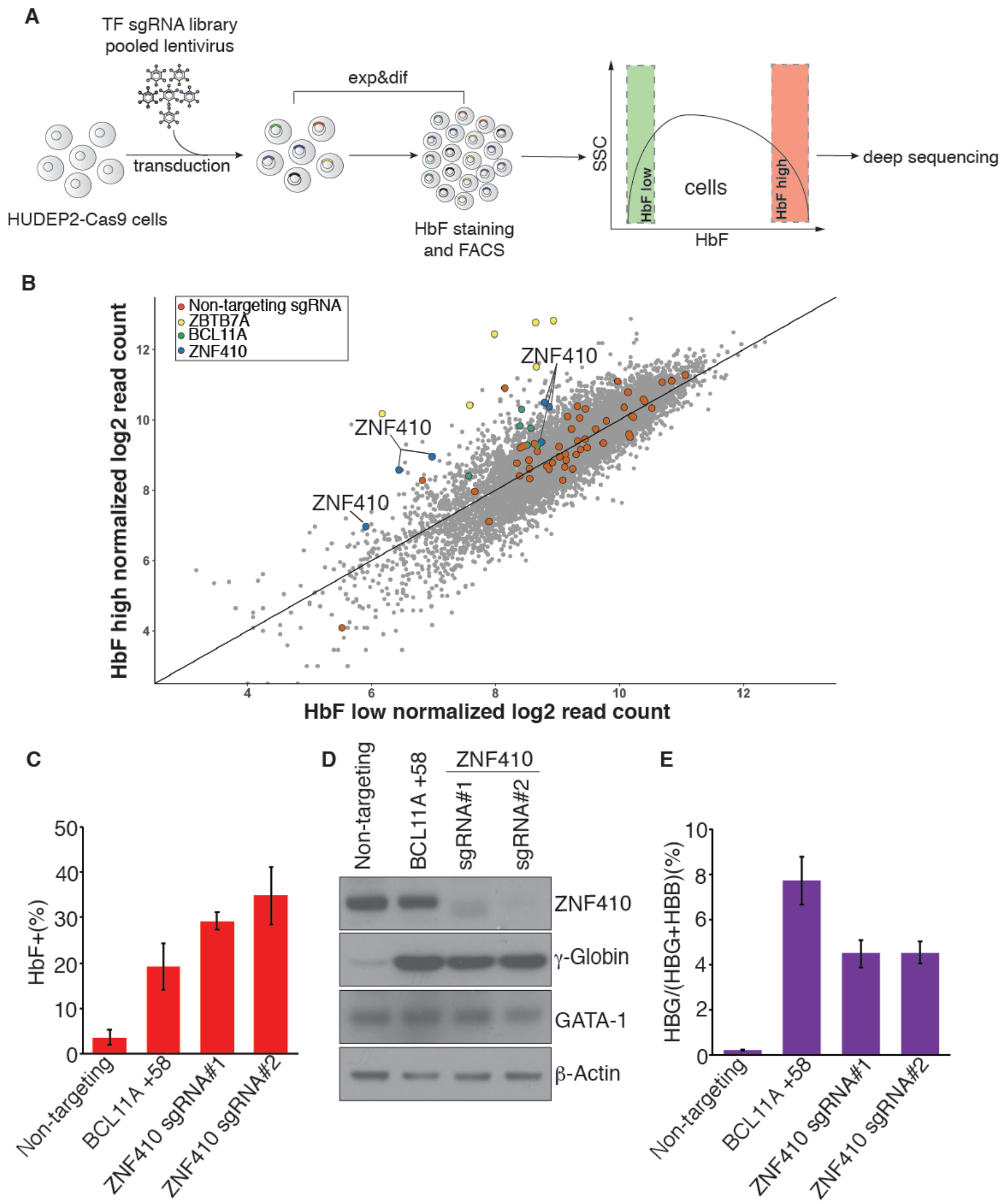
Domain-focused CRISPR-Cas9 screen identifies ZNF410 as a novel γ-globin repressor. (A) Schematic of screening strategy. TF: transcription factor. Exp&dif: expansion and differentiation. FACS: fluorescence-activated cell sorting. (B) Scatter plot of the screen results. Each dot represents one sgRNA. Control sgRNAs (red dots) are scattered randomly across the diagonal. ZBTB7A (yellow) and BCL11A (green) represent positive control sgRNAs. (C) Summary of HbF flow cytometric analyses. BCL11A +58: sgRNA targeting the +58 kb erythroid enhancer of the BCL11A gene serves as positive control. Non-targeting sgRNA serves as negative control. Results are shown as mean ± SD (n=3). (D) Immunoblot analysis using whole-cell lysates from differentiated HUDEP-2 cell pools transduced with indicated sgRNAs. (E) γ-globin mRNA measured by RT-qPCR in differentiated HUDEP-2 cells; data are plotted as percentage of γ-globin over γ-globin+ β-globin levels. Results are shown as mean ± SD (n=3).

Little is known about *ZNF410*. One report suggests that it functions as a transcriptional activator in human fibroblasts (Benanti et al., 2002). ZNF410 is widely expressed across human tissues (Genotype-Tissue Expression database). In blood, ZNF410 is highly expressed in the erythroid lineage (BloodSpot), and its mRNA levels are similar between fetal and adult erythroblasts (Huang et al., 2017).

To validate the screening results, two independent sgRNAs targeting the DNA-binding domain of ZNF410 were stably introduced into HUDEP-2-Cas9 cells along with a positive control sgRNA (targeting the +58 erythroid enhancer of the *BCL11A* gene) and non-targeting negative control sgRNA. Depletion of ZNF410 strongly increased the fraction of HbF-expressing cells as determined by flow cytometry using anti-HbF antibodies (Figures 1C and S1A). Western blotting revealed substantial elevation of γ-globin protein in ZNF410 depleted HUDEP-2 cells. Protein levels of GATA1 were unchanged, consistent with erythroid maturation being intact in these cells (Figure 1D). To assess whether ZFN410 impacts the transcription of the γ-globin gene, we carried out RT-qPCR. A robust increase in primary and mature γ-globin mRNA occurred upon ZFN410 depletion, suggesting transcriptional regulation (Figures 1E and S1B-S1C). ZNF410 loss did not impact ε-globin mRNA levels, suggesting specificity for the fetal globin genes (Figure S1D). Importantly, there were no notable changes in α-globin, β-globin, and GATA1 mRNA levels (Figures S1E-S1G), suggesting that ZNF410 depletion did not overtly impair erythroid differentiation, an observation further supported by RNA-seq analysis (below, Figure S1H). Taken together, our screen identified ZNF410 as novel repressor of γ-globin gene expression in HUDEP-2 cells.

### Depletion of ZNF410 elevates γ-globin levels in primary human erythroblasts

The repressive role of ZNF410 on HbF was further tested in primary human erythroblasts derived from a three-phase human CD34+ hematopoietic stem and progenitor cells (HSPCs) culture system (Grevet et al., 2018). We depleted ZNF410 by electroporating ribonucleoprotein (RNP) Cas9:sgRNAs complexes using two independent sgRNAs. A sgRNA targeting the erythroid +58 enhancer of BCL11A was used as positive control. In line with findings in HUDEP-2 cells, ZNF410 depletion significantly elevated the proportion of HbF+ cells (Figures 2A and 2B), γ-globin protein levels (Western blot, Figure 2C), HbF protein levels (HPLC, Figure 2D), and γ-globin primary and mature mRNA (Figures 2E and S2A-S2B). Moreover, ε-globin mRNA levels were unaffected (Figure S2C); α-globin, β-globin, and GATA1 levels remained largely unchanged (Figures S2D-S2F), suggesting that ZNF410 loss did not adversely affect maturation of these cells. Cell surface marker phenotyping using anti-CD71 and anti-CD235a antibodies as well as examination of cell morphology (Figures S2G and S2H) indicated normal erythroid maturation in ZNF410 deficient cells.

**Figure 2.**
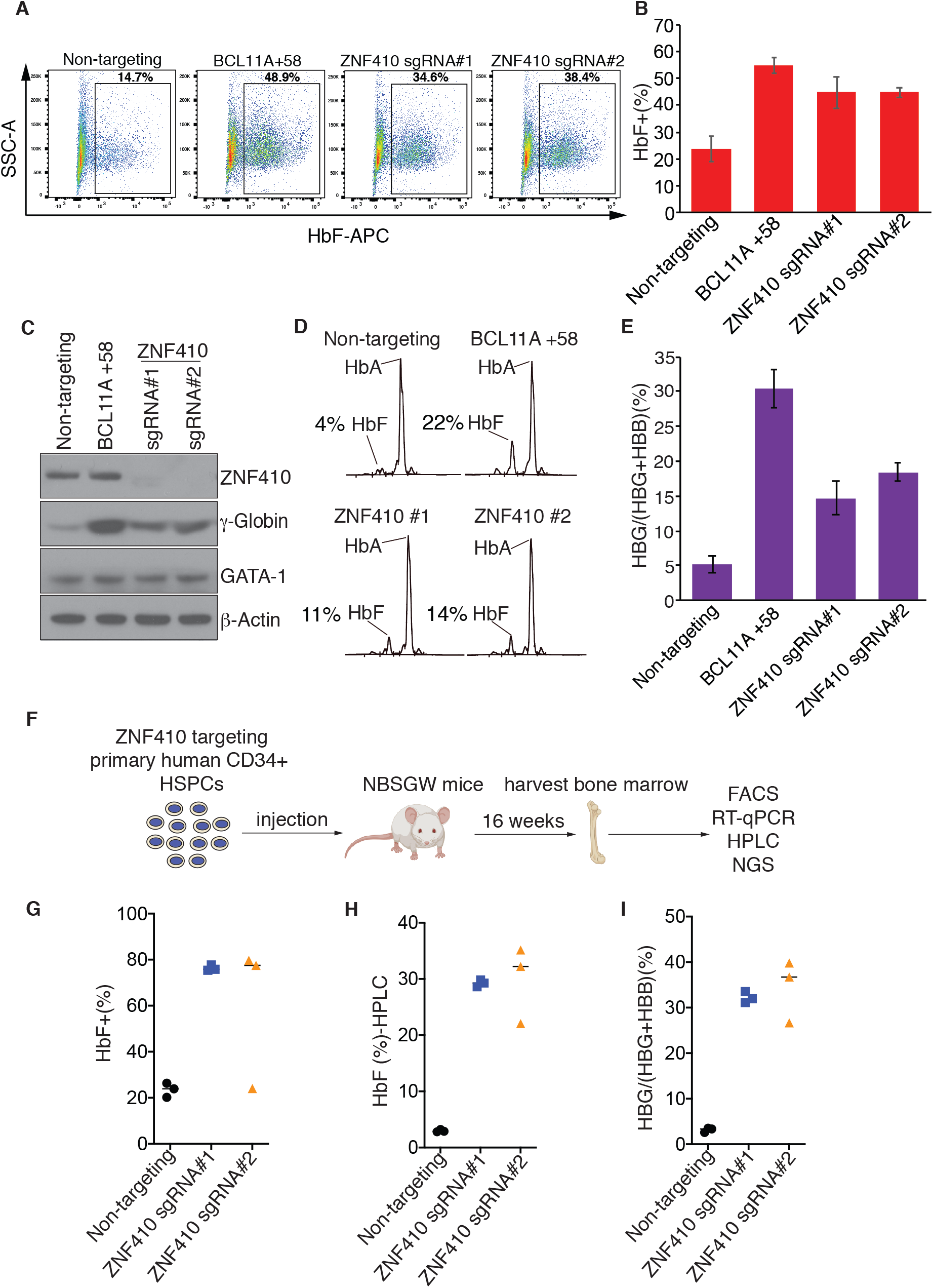
ZNF410 depletion induces γ-globin expression in primary erythroblasts. (A) Representative flow cytometric analysis of cells stained with anti-HbF antibody on day 15 of erythroid differentiation. BCL11A +58: sgRNA targeting the +58 kb erythroid enhancer of the BCL11A gene. (B) Summary of HbF flow cytometric analyses. Results are shown as mean ± SD (n=3 donors). (C) Immunoblot analysis using whole-cell lysates from primary erythroblasts with indicated sgRNAs on day 15 of differentiation. (D) Representative HPLC analysis of cells with indicated sgRNAs on day 15 of differentiation. HbA: hemoglobin A (adult form); HbF: fetal hemoglobin. HbF peak area is showed as percent of total HbF+HbA. (E) γ-globin mRNA measured by RT-qPCR in primary erythroblasts on day 12 of differentiation; data are plotted as percentage of γ-globin over γ-globin+ β-globin levels. Results are shown as mean ± SD (n=3 donors). (F) Schematic of experimental design. (G-I) Summary of HbF flow cytometric analyses (G), HPLC analysis (H) and γ-globin mRNA measured by RT-qPCR (I) in human CD235a+ erythroblasts isolated from recipient bone marrows. Each dot represents a single recipient mouse. n=3 mice per sgRNA.

In vitro cell culture systems may not always reflect the true biology of developing erythroid cells in vivo. Moreover, commonly used mouse models may in some cases not faithfully reproduce all regulatory features of human erythroid cells (Huang et al., 2020). Therefore, we used a human-to-mouse xenotransplantation model to further assess the role of ZNF410 on the regulation of γ-globin in vivo. We transfected normal adult human donor CD34+ HSPCs with ribonucleoprotein complex consisting of spCas9 + two sgRNAs (analyzed separately) or a non-targeting sgRNA as negative control, and then transplanted them into NBSGW immunodeficient mice that support human erythropoiesis in the bone marrow (McIntosh et al., 2015). We measured the fraction of various engrafted human lineages and their gene editing frequencies in recipient bone marrow at 16 weeks after xenotransplantation (Figure 2F), a time at which CD34+ progenitor cells are mainly derived from the human transplant (McIntosh et al., 2015). Donor chimerism of ZNF410 edited CD45+ hematopoietic cells was slightly lower than in control cells exposed to non-targeting sgRNA (Figure S2I). Chimerism levels and indel frequencies were similar in all edited and nonedited lineages tested, including B-cells, myeloid, erythroid and progenitor cells (Figures S2J and S2K), indicating that ZNF410 depletion did not overtly impact hematopoietic development. Importantly, in the erythroid compartment (CD235+), we observed a robust increase in the fraction of HbF+ cells (from ∼23% to ∼78%), HbF protein levels (from ∼3% to ∼33%), and γ-globin mRNA levels (from ∼3% to ∼32%) (Figures 2G-2I), consistent with the results in HUDEP-2 and cultured primary human erythroblasts. Again, depletion of ZNF410 did not appear to impair erythroid maturation (Figure S2L). Collectively, these in vivo studies verify that ZNF410 functions as a robust repressor of HbF with little effect on hematopoietic development.

### ZNF410 represses HbF by modulating CHD4 expression

ZNF410 was previously reported to function as a transcriptional activator (Benanti et al., 2002). To understand how ZNF410 regulates the transcription of γ-globin genes, we performed RNA-seq experiments in ZNF410 depleted differentiated HUDEP-2 cells and primary human erythroblasts. Upon ZNF410 depletion in HUDEP2 cells, 70 genes were up- and 46 genes were down-regulated, respectively, with a threshold setting of 1.5-fold (p-value<0.05), and only counting genes that incurred changes with both ZNF410 sgRNAs in each of the biological replicates (Figure S3A). In primary erythroid cultures, 83 genes were up-regulated and 126 genes were down-regulated, respectively upon ZNF410 depletion (Figure S3A). This includes 30 up-regulated and 15 down-regulated genes in both cell types (Figures S3B). Notably, γ-globin (HBG) mRNA levels stood out among the most strongly induced genes (Figures 3A-3B and S3C-S3D, Table S8).

**Figure 3.**
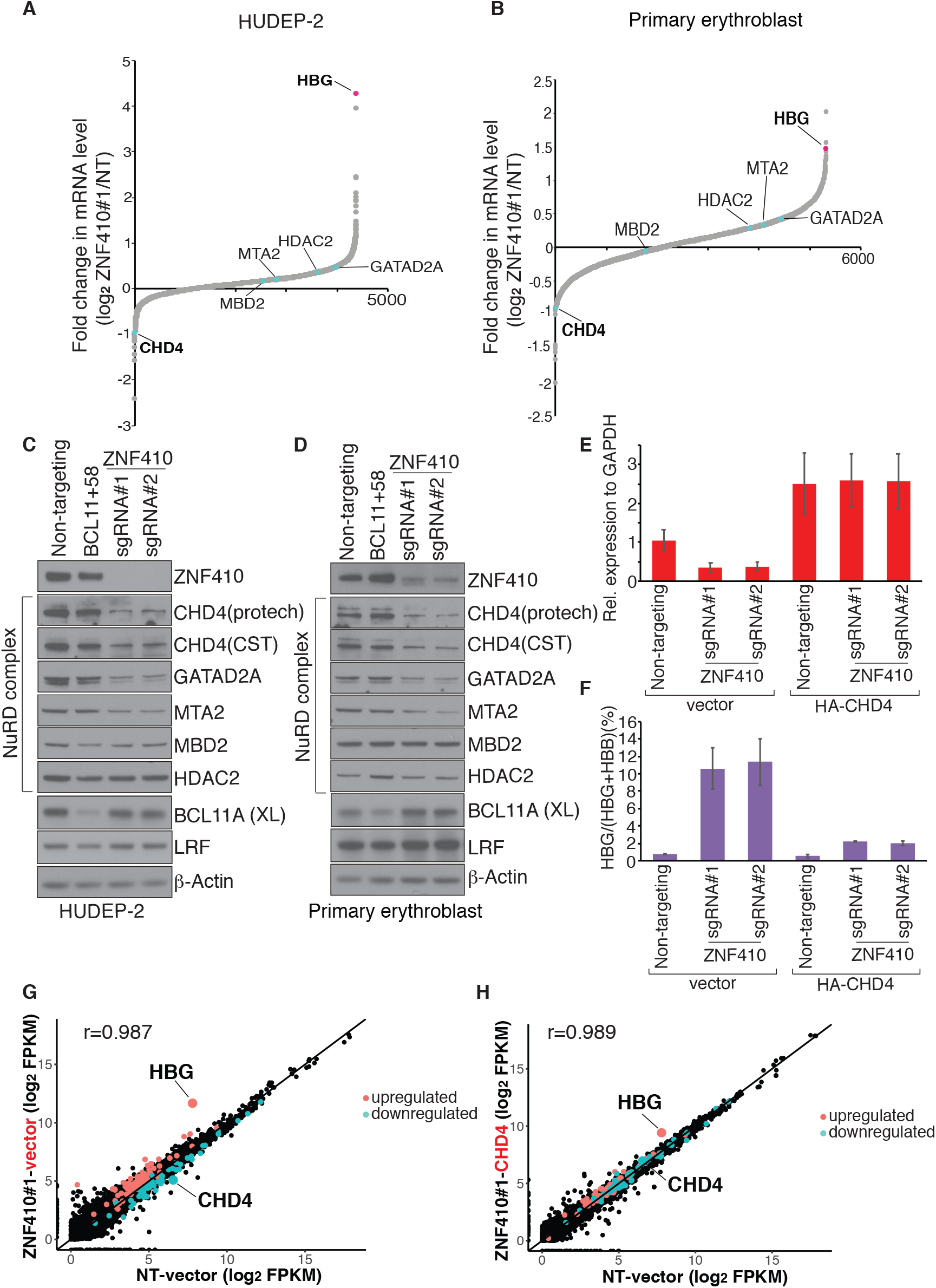
CHD4 mediates γ-globin repression by ZNF410. (A) RNA-seq analysis of HUDEP-2 cells transduced with ZNF410 sgRNA#1. Infected cells were sorted and differentiated for 7 days. Plotted is the average fold-change in mRNA levels of two biological replicates. Genes encoding NuRD complex subunits and γ-globin (HBG) are indicated. Fragments per kilobase of transcript per million (FPKM) mapped reads were used to calculate fold change. NT: non-targeting. (B) RNA-seq analysis of primary erythroblasts with ZNF410 depletion by sgRNA#1. Cells were differentiated for 12 days. Plotted is the average fold-change in mRNA levels of two independent donors. (C-D)Immunoblot analysis using whole-cell lysates from differentiated HUDEP-2 cells (C) and primary erythroblasts on day 15 of differentiation (D). BCL11A(XL) is the functional BCL11A isoform. CHD4 antibodies were from Proteintech and Cell Signaling Technologies (CST). (E) CHD4 mRNA levels measured by RT-qPCR in ZNF410 deficient HUDEP-2 cells transduced with lentiviral vector containing CHD4 cDNA or empty vector. Results are shown as mean ± SD (n=2). GAPDH was used for normalization. (F) γ-globin levels measured by RT-qPCR in ZNF410 deficient HUDEP-2 cells transduced with lentiviral vector containing CHD4 cDNA or empty vector, data are plotted as percentage of γ-globin over γ-globin+ β-globin levels. Results are shown as mean ± SD (n=2). (G) Scatter plot of RNA-seq analysis in ZNF410 deficient HUDEP-2 cells (by ZNF410 sgRNA#1) with empty vector. Cells with non-targeting sgRNA and vector serve as control. Each dot indicates a gene. Each gene is depicted according to averaged FPKM value from 2 biological replicates. r: Pearson’s correlation coefficient. NT: non-targeting. (H) Scatter plot of RNA-seq analysis in ZNF410 deficient HUDEP-2 cells (by ZNF410 sgRNA#1) with re-introduction of CHD4 cDNA. Each gene is depicted according to averaged FPKM value from 2 biological replicates. r: Pearson’s correlation coefficient.

The CHD4 gene which encodes the catalytic subunit of the NuRD complex, was among the most downregulated genes (Figures 3A-3B and S3C-S3D, Table S8). The NuRD complex contributes to the γ-globin repressive functions of BCL11A and LRF in erythroid cells. In addition to CHD4, the GATAD2A, HDAC2, MBD2 and MTA2 subunits of the NuRD complex are required for γ-globin repression (Sher et al., 2019). However, the mRNA levels of these subunits were not diminished in ZNF410 depleted cells, suggesting that CHD4 is the only ZNF410-regulated NuRD subunit (Figures 3A-3B and S3C-S3D). We validated these results by RT-qPCR in HUDEP-2 cells, cultured primary erythroid cells, and ZNF410-depleted erythroid cells isolated from xenotransplanted NBSGW mice (Figures S3E-S3G). Overall, the reduction in CHD4 transcript levels amounted to approximately 65% in HUDEP-2, primary cultures and xenotransplanted mice (Figures S3E-S3G). That ZNF410 might be limiting for CHD4 transcription is further supported by the strong correlation between ZNF410 and CHD4 transcript levels across 53 human tissues based on the Genotype-Tissue Expression database (Figure S3H).

In agreement with the mRNA analysis, CHD4 protein levels were significantly reduced upon ZNF410 depletion in HUDEP2 and primary erythroblasts, while BCL11A, LRF, HDAC2 and MBD2 protein amounts remained unchanged (Figures 3C and 3D). Of note, although GATAD2A and MTA2 transcripts were unaltered, their proteins levels were reduced upon ZNF410 depletion (Figures 3C-3D and S3E-S3G). We speculate that these subunits are destabilized in the absence of CHD4 (Torrado et al., 2017). No other genes known to regulate γ-globin silencing were altered by ZNF410 depletion, suggesting that CHD4 is the critical link between ZNF410 and γ-globin silencing.

### CHD4 is the sole mediator of ZNF410 function

Our results so far implicate CHD4 as the key ZNF410-controlled regulator of γ-globin silencing. Therefore, we examined whether expression of CHD4 in ZNF410 depleted cells restored γ-globin silencing. We transduced ZNF410-deficient HUDEP-2 cells with a lentiviral vector encoding *CHD4* cDNA linked to an IRES element and GFP expression cassette, followed by FACS purification of GFP+ cells. The transduced cells expressed *CHD4* mRNA at a level approximately 2.0-fold above normal (Figure 3E). CHD4 expression almost completely restored the silencing of γ-globin without influencing the expression of other erythroid genes such as α-globin, β-globin, and GATA1 (Figures 3F and S4A-S4E). To assess whether other transcriptional changes resulting from ZNF410 loss are also attributable to lower CHD4 levels, we performed replicate RNA-seq experiments in the CHD4-expressing ZNF410-deficient HUDEP-2 cells. Notably, 69 out of 70 upregulated genes and 44 out of 46 downregulated genes in the ZNF410 deficient cells were expressed at normal levels following CHD4 “rescue” (Figures 3G-3H and S4F-S4G). Why the expression of three genes was incompletely restored upon *CHD4* re-expression is unclear but might be due to imperfect levels of CHD4 restoration or a drift in gene expression profiles following gene knockout/rescue experiments in cell pools. None of the three genes whose expression remained unrestored to normal levels are associated with ZNF410 ChIP-seq peaks (see below), suggesting that they are not direct ZNF410 targets. Together, these results suggest that CHD4 is the only functionally relevant ZNF410 target gene, and is responsible for the repression of γ-globin transcription. Another remarkable finding is that the γ-globin genes (HBG1/2) are among the most sensitive to CHD4 levels.

### A singular enrichment of ZNF410 binding clusters at the CHD4 gene

Our RNA-seq study identified numerous genes that were up- or down-regulated after ZF410 depletion. To investigate which of these genes are direct ZF410 targets, we performed anti-ZNF410 ChIP-seq in HUDEP-2 and primary human erythroblasts with ZNF410-deficient HUDEP-2 cells as a control. We detected only 8 high-confidence peaks corresponding to 7 genes total in both HUDEP-2 and primary human erythroid cells (Figure 4A). To exclude the possibility that such unusually few called peaks are due to limitations in ZNF410 detection by ChIP, we overexpressed HA-tagged ZNF410 or empty vector in HUDEP-2 cells (Figure S5A). Anti-HA ChIP-seq detected the same 8 ZNF410 peaks with comparable intensity profiles (Figure 4A). No ZNF410 binding was detected at the β-globin locus (Figure S5B), supporting a model in which ZNF410 regulates γ-globin transcription indirectly. 6 of 8 ZNF410 ChIP-seq peaks were of modest magnitude. Most strikingly, two very strong peaks were located at the promoter and enhancer of the CHD4 gene. These data suggest that ZNF410 directly regulates an unusually small number of genes and that suppression of ZNF410 may induce γ-globin transcription by downregulating the NuRD component CHD4.

**Figure 4.**
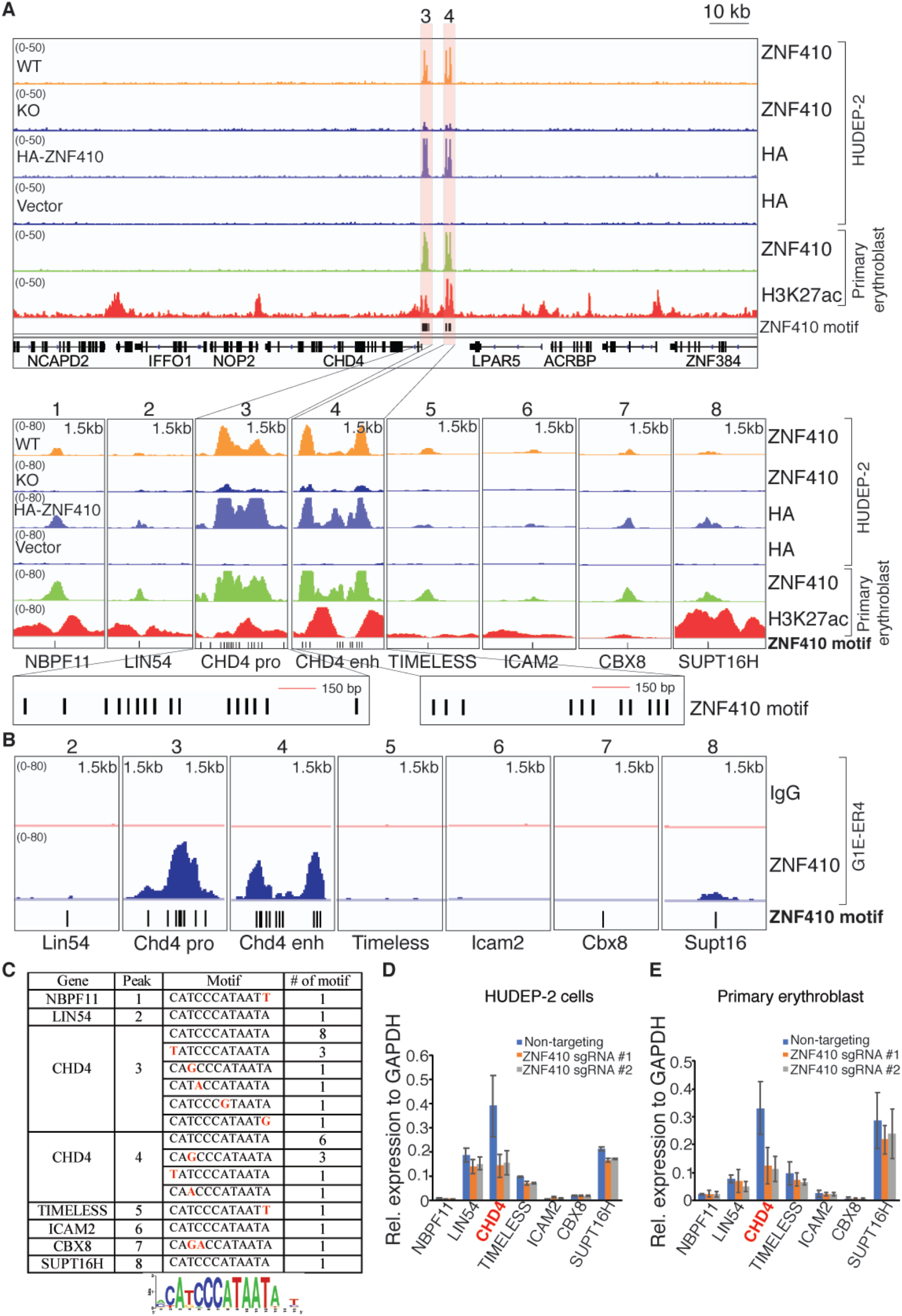
ZNF410 binding to the CHD4 locus occurs at highly conserved motif clusters. (A) ChIP-seq profiles of endogenous ZNF410, HA-ZNF410 and H3K27ac. CHD4 promoter and enhancer are highlighted in orange. ZNF410 binding motifs are denoted by vertical black lines at the bottom. The 8 peak-associated genes are shown below the tracks. ZNF410 KO cells and cells transduced with empty vector serve as negative controls. HA-ZNF410: N-terminal HA tagged ZNF410. HA: hemagglutinin. (B) Browser tracks of endogenous ZNF410 ChIP-seq occupancy at the 7 murine counterparts in differentiated mouse erythroid cells. ZNF410 binding motifs are showed at the bottom. IgG track serve as negative control. (C) Summary of ZNF410 binding motif counts at the 8 peaks, and derived de novo motif logo in the human genome. Red font indicates the sequence variants. (D-E) mRNA levels of the 7 ZNF410 bound genes in HUDEP-2 cells transduced with indicated sgRNAs (D) and primary erythroblasts electroporated with indicated sgRNAs (E) by RT-qPCR (n=2). GAPDH was used for normalization.

HOMER motif analysis based on the 8 high-confidence binding sites from our ChIP-seq data generated the 12-nucleotide motif CATCCCATAATA (Figure 4C), which is almost identical to that found by in vitro SELEX experiments of human ZNF410 (Jolma et al., 2013). Utilizing EMBOSS fuzznuc we found 434 of such motif instances, and 3677 instances when combining all the motifs found under ZNF410 ChIP-seq peaks. Overall frequency of these motifs is very low compared to those of most transcription factor binding sites (Srivastava and Mahony, 2020). However, since the vast majority of these motifs had no measurable ChIP-seq signal, additional features must account for the rare in vivo binding events. The two strongest ZNF410 ChIP-seq peaks at the CHD4 promoter and enhancer encompass 15 and 11 motifs, respectively (each within a 1.5 kb window), while the remaining 6 modest peaks harbor only one motif (Figures 4A and 4C). Importantly, the 2 peaks at the CHD4 locus are the only regions in the entire genome with a high density of ZNF410 motifs, likely explaining the exquisite target specificity.

To explore additional criteria that might account for the selectivity of ZNF410 binding to chromatin, we asked whether ZNF410 chromatin occupancy is associated with features of open chromatin. First, we generated in primary human erythroblasts ChIP-seq profiles for H3K27ac, a histone mark associated with active regulatory elements, and complemented these data by mining chromatin accessibility (ATAC-seq) data from primary human erythroblasts (Ludwig et al., 2019). All 8 ZNF410 peaks, including the two strong peaks at the CHD4 promoter and enhancer, fell into accessible chromatin that was flanked by regions enriched in H3K27ac (Figures 4A and S5C). In contrast, the vast majority of the unbound consensus motif instances elsewhere in the genome were in regions devoid of H3K27ac or ATAC-seq signal (Table S9). Although the scarcity of ZNF410-bound sites precludes an extensive correlative analysis, we did find only a very modest positive correlation between strengths of signal for ZNF410 binding with H3K27ac levels or ATAC-seq signal (Pearson’s correlation coefficient of 0.33 and 0.24, respectively in primary erythroblasts; Figure S5C). However, the categorical association of ZNF410 binding with open, active chromatin was almost complete, suggesting that ZNF410 requires open chromatin and perhaps additional transcription factor binding sites, in addition to motif clustering, to enable its binding to chromatin.

The mere occupancy of a transcription factor at a gene does not necessarily lead to regulatory influence. We examined and validated our RNA-seq data in ZNF410-deficient cells for the expression of the seven genes bound by ZNF410. Importantly, among these genes, *CHD4* was the only one with significantly reduced mRNA levels in ZNF410 depleted cells (Figures 4D-4E and S5D-S5E). Thus, ZNF410 directly and functionally regulates a single target gene, CHD4, in erythroid cells. Hence, the vast majority of gene expression changes that occur upon ZNF410 loss are likely due to diminished CHD4 levels. This model is supported by the restoration of transcriptome changes upon CHD4 expression in ZNF410 deficient cells (Figures 3G-3H and S4F-S4G).

### ZNF410 binding to chromatin occurs at highly conserved motif clusters

Highly conserved non-coding elements can function as enhancers and are associated with transcription factor binding sites (Pennacchio et al., 2006). We assessed conservation of the ZNF410 binding regions at the CHD4 locus using the phastCons scores deduced from sequence similarities across 100 vertebrate species (Siepel et al., 2005). Both ZNF410 binding site clusters displayed a high degree of conservation, comparable to those at the *CHD4* exons (Figure S5F). Moreover, the human ZNF410 protein sequence is 94% identical to mouse protein, and the DNA binding ZF domain is nearly 100% identical (Figure S5G).

To examine whether ZNF410 binding selectivity for the CHD4 locus is conserved in mouse, we carried out ZNF410 ChIP-seq in the erythroid cell line G1E-ER4 (Weiss et al., 1997). As in human cells, the Chd4 promoter and enhancer were by far the most strongly occupied regions genome wide (Figure 4B). Of the 6 human genes that exhibited modest ZNF410 ChIP-seq signals, no signal was detected at 4 orthologues in mice (*Lin54, Timeless, Icam2*, and *Cbx8*). A modest signal was detected at the mouse *Supt16* gene but its expression was not altered by loss of ZNF410. Further analysis confirmed that among 1876 motif instances matching the most common motifs in the human genome, the *Chd4* promoter and enhancer are also by far the most enriched locations in mouse genome (Figure 4B). Taken together, these findings indicate that regulation of the *CHD4* gene by ZNF410 is mediated through unique, evolutionarily conserved motif clusters.

### Characterization of DNA binding by ZNF410

ZNF410 contains five tandem C2H2-type zinc fingers (ZFs) potentially involved in DNA binding (Figure 5A). However, as is the case for many ZF transcription factors, not all ZFs necessarily make direct DNA contacts. We assessed direct DNA binding by full length (FL) ZNF410 to sequences found at the CHD4 gene by electrophoretic mobility shift assays (EMSAs). Using nuclear extracts from COS cells overexpressing FLAG-tagged ZNF410 constructs and radiolabeled probes containing the relevant motifs (Figure 5B), FL ZNF410 protein displayed comparable binding to each of the four DNA probes containing a single motif associated with ChIP-seq signal at the CHD4 promoter and enhancer (Figure 5C). Addition of anti-FLAG antibody to the binding reaction led to a “supershift” (Figure 5C), confirming binding specificity.

**Figure 5.**
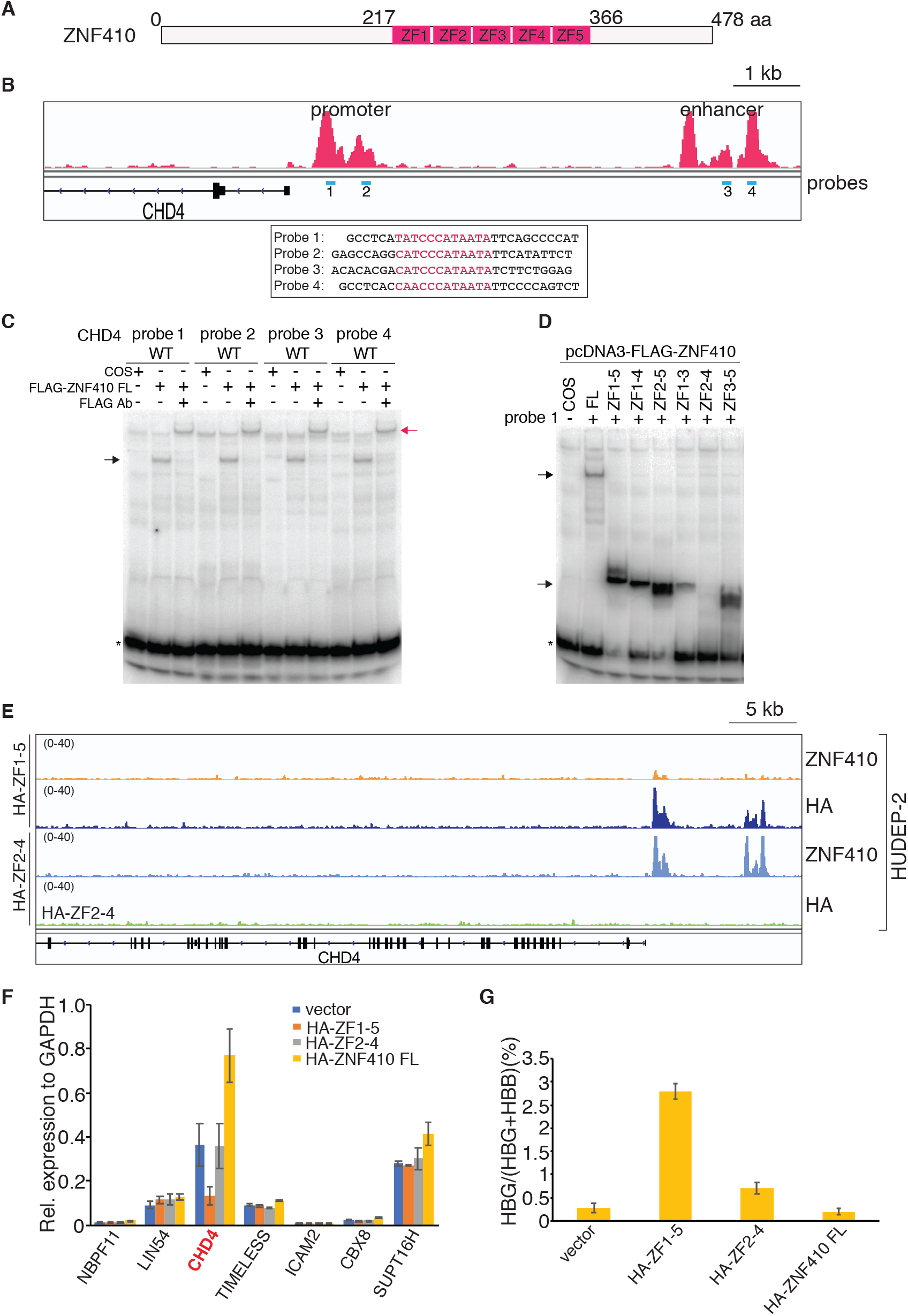
The ZF domain of ZNF410 is sufficient for DNA binding in vitro and in vivo. (A) Schematic of human ZNF410. ZNF410 contains 478 amino acids, with five C2H2-type zinc fingers (ZF) between amino acids 217 and 366. (B) ZNF410 ChIP-seq track with EMSA probes shown in blue underneath the peaks and probe sequences showed below. Motifs are highlighted in red. (C) Full-length ZNF410 binds to the four motifs from the CHD4 promoter and enhancer sites. Black arrow: ZNF410-probe complex; red arrow: FLAG antibody-ZNF410-probe complex. *: free probes. Ab: antibody, FL: full length. FLAG-ZNF410: N-terminal FLAG tagged ZNF410. (D) The ZF domain of ZNF410 and its truncations bind to the motif except ZF2-4. Black arrow: i ZNF410 FL, ZF domain or domain truncation-probe complex. *: free probes. (E) Browers track of endogenous ZNF410, HA ChIP-seq occupancy at the CHD4 locus in HUDEP-2 cells overexpressing HA-ZF1-5 or HA-ZF2-4. HA: hemagglutinin. (F) mRNA levels of the 7 ZNF410 bound genes by RT-qPCR in differentiated HUDEP-2 cells with HA-ZF1-5, HA-ZF2-4 or HA-ZF410 FL overexpression. Results are shown as mean ± SD (n=2). GAPDH was used for normalization. FL: full length. (G) γ-globin levels measured by RT-qPCR in differentiated HUDEP-2 cells with HA-ZF1-5, HA-ZF2-4 or HA-ZF410 FL overexpression. Data are plotted as percentage of γ-globin over γ-globin+ β-globin levels. Results are shown as mean ± SD (n=2).

The domain spanning all five ZFs was sufficient for DNA binding, and like FL ZNF410, displayed similar binding intensity across the four probes (Figure S6A). Again, addition of anti-FLAG antibody caused a “supershift”, validating the specific interactions between the ZF domain and the probes (Figure S6A). The stronger signal generated by the ZF domain compared to that of full length ZNF410 (Figure 5D) is likely due to the higher expression level of the former (Figure S6B). To assess the contribution to DNA binding by each of the five ZFs, we generated versions containing various ZF combinations (Figure S6B). In EMSA, the central 3 ZFs (2-4) were insufficient for DNA binding, however, if either ZF1 or ZF5 was present (ZF1-4 and ZF2-5, respectively), DNA binding was enabled (Figure 5D). Additionally, we observed DNA binding activity, albeit reduced, by ZF1-3 and ZF3-5 (Figure 5D). Thus, the central 3 ZFs (2-4) are insufficient for DNA binding, with a ZF at either end (ZF1 or ZF5) contributing to DNA contacts in this assay. Together, these data support the view that each of the five ZFs of ZNF410 is involved in DNA binding.

According to the EMSAs, the ZF1-5 domain displays strong DNA binding in vitro, while ZF2-4 does not (Figure 5D). We reasoned that overexpression of ZF1-5 but not ZF2-4 should compete with endogenous ZNF410 for chromatin binding, thus acting in a dominant-negative manner. To test this hypothesis, we introduced via lentiviral infection into HUDEP-2 cells a construct containing HA-tagged ZF1-5 or ZF2-4 driven by the EF1α promoter. As control, we also forced the expression of full-length HA-tagged ZNF410. Overexpression of ZF1-5 or ZF2-4 did not influence the endogenous ZNF410 expression (Figure S6C). ChIP-seq experiments demonstrated that overexpressed ZF1-5 (roughly 20-fold compared to endogenous ZNF410 protein levels) bound to the CHD4 regulatory regions and to the other ZNF410 targets in a pattern very similar to endogenous ZNF410 (Figures 5E and S6C-S6E). Moreover, full-length HA-ZNF410 when overexpressed to similar levels to HA-ZF1-5 (Figure S6C) also produced similar binding patterns (Figure 4A). This suggests that the ZF domain is sufficient for ZNF410 chromatin occupancy, and that regions outside this domain contribute little if anything to chromatin binding. ZF2-4 displayed no chromatin occupancy at all sites examined, consistent with in vitro DNA binding properties, but with the caveat that it was expressed at lower levels (Figure 5E and S6C-S6E). Accordingly, ZF1-5 markedly interfered with endogenous ZNF410 binding while ZF2-4 was inert (Figure 5E and S6D-S6F). To assess the impact of interference with endogenous ZNF410 chromatin binding on CHD4 expression, we carried out RT-qPCR and found CHD4 mRNA levels to be reduced by approximately 70% compared to control (Figure 5F), which is comparable to ZNF410 knockout (Figure 3E). In contrast, overexpression of ZF2-4 did not influence CHD4 expression, however, overexpression of full length ZNF410 increased CHD4 expression (Figure 5F). Thus, the ZF region is sufficient for chromatin occupancy but insufficient for *CHD4* gene activation. In agreement with the results from ZNF410 depletion experiments, overexpression of ZF1-5 or ZF2-4 did not impact expression of the other 6 ZNF410 bound genes (Figure 5F). Finally, we also measured the impact of ZNF410 expression on γ-globin levels. ZF1-5 expression triggered a significant increase in γ-globin mRNA levels (Figures 5G and S6G), again comparable to that observed in ZNF410 depleted cells, while ZNF410 FL expression led to a slight decrease in the already low γ-globin mRNA levels (Figure 5G and S6G). We did note a modest increase in γ-globin mRNA levels upon ZF2-4 expression, the reason for which is unknown. In sum, the ZF domain of ZNF410 is necessary and sufficient to bind to DNA in vitro and in vivo and does not seem to bear any transactivation function on its own.

### Structural basis of ZNF410-DNA binding

To further gain insight into the molecular basis of how the ZNF410 tandem ZF domain recognizes its targeting DNA sequence, we performed crystallization of the ZNF410-DNA complex. We first quantified the binding affinity of the ZNF410 ZF domain (ZF1-5) with the consensus motif by fluorescence polarization using purified GST fusion protein (Patel et al., 2016a). The ZF domain displayed a dissociation constant (K_D_) of 22 nM for the oligo containing the consensus motif while there was no measurable binding to the negative control (that shares 7/17 bp with the consensus motif) under the same conditions (Figure 6A). Using the same samples, we confirmed that the binding affinity of the ZF domain with the consensus oligo was between 10 and 20 nM by EMSA (Figure S6H). Next, we determined the crystal structure of the ZF domain in complex with the same 17-bp oligo containing the consensus motif. The structure of the protein-DNA complex was solved by the single-wavelength anomalous diffraction (SAD) method (Hendrickson et al., 1990) at 2.75 Å resolution (Table S10). As in conventional C2H2 ZF proteins (Wolfe et al., 2000), each of five fingers of ZNF410 comprises two strands and a helix, with two histidine residues in the helix together with one cysteine in each strand coordinating a zinc ion, forming a characteristic tetrahedral C2-Zn-H2 structural unit that confers rigidity to the fingers. When bound to DNA, ZF1-5 occupies the DNA major groove, with their α-helices toward DNA and the strands and the C2-Zn-H2 units facing outside (Figures 6B and C). Side chains from specific amino acids within the N-terminal portion of each helix and the preceding loop (i.e., the 7 residues prior to the first Zn-coordinating histidine; Figure 6D) make major groove contacts with primarily three adjacent DNA base pairs, which we term the “triplet element”. The DNA oligo used for crystallization contains the 15-bp consensus sequence (numbered 1-15 from 5’ to 3’ of the top strand; colored magenta in Figure 6E) recognized by the five fingers, plus one additional base pair on each end of the DNA duplex. The protein sequence runs in the opposite direction of the top strand, from carboxyl (COOH) to amino (NH2) termini, resulting in ZF5 recognizing the 5’ triplet (bp position 1-3), and the ZF1 recognizing the 3’ triplet (bp position 13-15) (Figure 6E).

**Figure 6.**
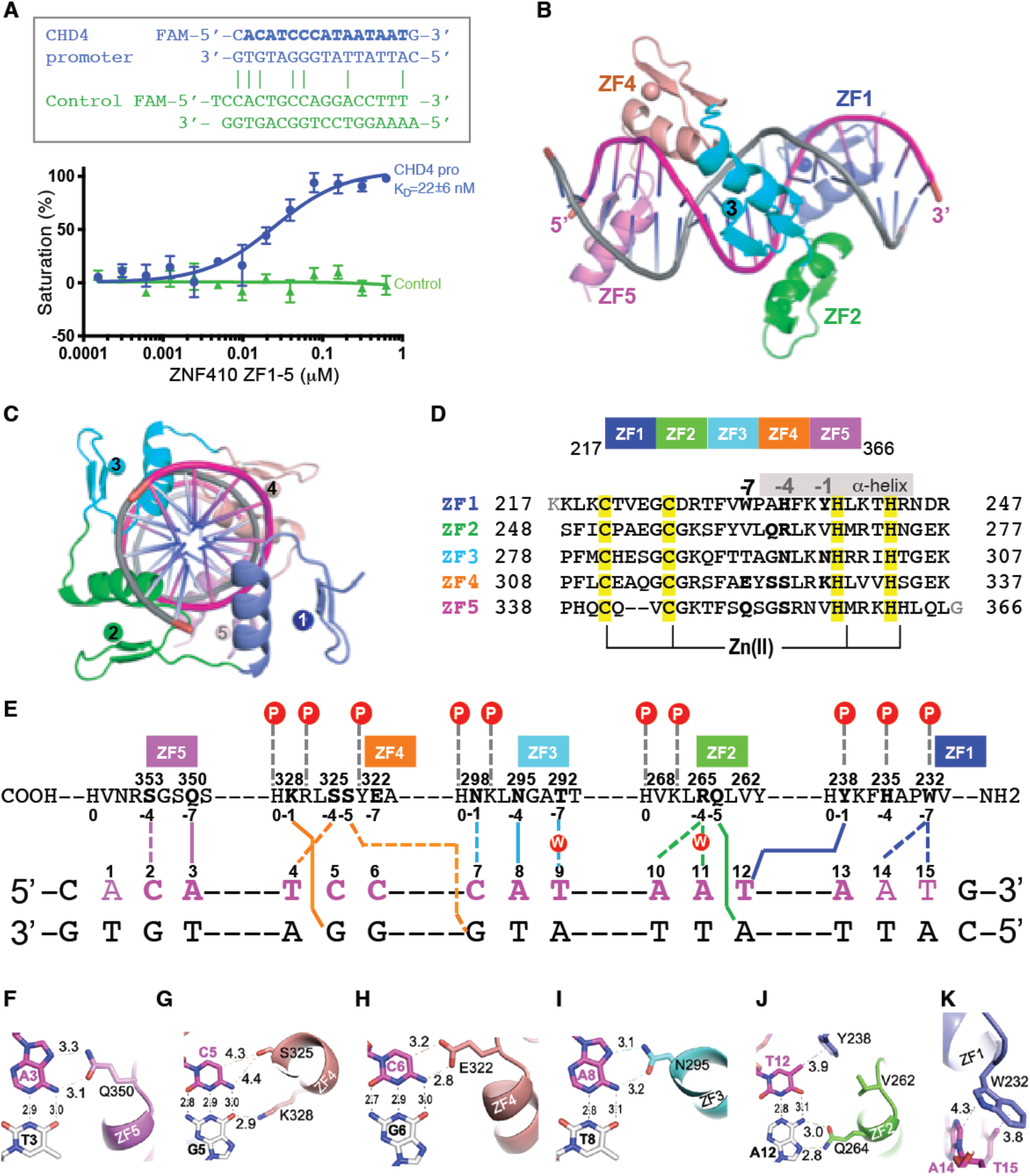
Structural basis of ZNF410-DNA binding. (A)Binding affinity measurements of the ZF domain against oligos by fluorescence polarization assays. Above: oligos used. X-axis: concentration of purified GST-ZF1-5 protein, Y-axis: percentage of saturation. Pro: promoter. (B-C)Two ortholog views of a ZNF410 ZF1-5 binding to DNA. (D) Sequence alignment of the five zinc fingers of ZNF410 with DNA base-interacting positions −1, −4 and −7 in bold. The Zn-coordinating residues C2H2 of each finger are highlighted in yellow. (E) General scheme of interactions between ZF1-5 and DNA. The top line indicates amino acids of each finger from C to N terminus. The first zinc-coordination His in each finger is referenced as position 0, with residues before this, at sequence positions −1, −4 and −7, corresponding to the 5’-middle-3’ of each DNA triplet element. The bottom two lines indicate the sequence of the double-strand oligonucleotide used for crystallization. The base pair matching the consensus sequence by are numbered as 1-15. (F-K) Examples of base-specific contacts between each ZF and DNA. (F) Q350 of ZF5 interacts with A3. (G) S325 and K328 of ZF4 interacts with the C:G base pair at position 5. (H) E322 of ZF4 interacts with C6. (I) N295 of ZF3 interacts with A8. (J) Q264 of ZF2 and Y238 of ZF1 interact with the T:A base pair at position 12. (K) W232 of ZF1 interacts with A14 and T15.

Each zinc finger contributes to specific DNA interactions. The most dominant direct base-specific interactions observed are the Ade-Gln and Ade-Asn contacts via three fingers, e.g., Q350 of ZF5 interaction with A3 (Figure 6F), N295 of ZF3 interaction with A8 (Figure 6I), and Q264 of ZF2 interaction with A12 (Figure 6J). In accordance with apposition of Gln/Asn with Ade as the most common mechanism for Ade recognition (Luscombe et al., 2001), the side chain carboxamide moiety of glutamine and asparagine donates a H-bond to the *O7* and accepts a H-bond from the *N6* atoms of adenines, respectively, a pattern specific to Ade. ZF4 contacts two C:G base pairs at positions 5 and 6. K328 of ZF4 interacts with the *O6* atom of G5 (Figure 6G), while E322 of ZF4 forms a H-bond with the *N4* atom and a C-H•••O type H-bond (Horowitz and Trievel, 2012) with the *C5* atom of cytosine at position 6 (Figure 6H). In addition, S325 of ZF4 forms a van der Waals contact with cytosine at position 5 (Figure 6G). ZF1 uses two aromatic residues (Tyr238 and Trp232) for interaction with the methyl group of thymine at position 12 and 15, respectively (Figure 6J and 6K). Among the base specific interactions, the C:G base pair at position 5 and the T:A base pair at position 12 have direct protein interactions with both bases (Figure 6G and 6J). For additional description of the non-specific contacts, see supplementary text. In sum, the base specific interactions protect 10 base pairs (positions A3 to T12 in Figure 6E), out of 12-base pair consensus sequence (Figure 4C).

In addition to the direct base interactions, the first four fingers (ZF1-4) interact with DNA backbone phosphate groups, while ZF5 is devoid of such contact (Figure 6E). Among the five ZFs, ZF5 has the least number of contacts with DNA, whereas ZF1 has only the van der Waals contacts with the bases, and two of them are outside of consensus (nucleotide positions 14 and 15 in Figure 6E). This observation might explain that DNA binding in vitro was still enabled if either outside finger (ZF1 or ZF5), but not both, are removed (Figure 5D). Our structure also revealed that the fingers in the middle (ZF2-4) follow the one-finger-three base rule, each involving highly base specific interactions, whereas the fingers in the ends vary from 2-base (ZF5) to 4-base contacts (ZF1).

## Discussion

By leveraging an improved CRISPR-Cas9 screening platform, we identified ZNF410, a pentadactyl zinc finger protein, as a novel regulator of fetal hemoglobin expression. ZNF410 regulates γ-globin expression through selective activation of CHD4 transcription. CHD4 appears to be the only direct functional target of ZNF410 in erythroid cells. Two highly conserved clusters of ZNF410 binding sites at the CHD4 promoter proximal region and enhancer that appear to be unique in the human and mouse genomes account for selective accumulation of ZNF410 at the CHD4 locus. In the absence of ZNF410, CHD4 transcription is reduced but not entirely lost which explains the modest impact on global gene expression, and exposes the γ-globin genes as particularly sensitive to CHD4 levels. In vitro DNA binding assays and crystallography reveal the DNA binding modalities. This study thus illuminates a highly selective transcriptional pathway from ZNF410 to CHD4 to the γ-globin genes in erythroid cells.

Most transcription factors bind to thousands of genomic sites, of which a significant fraction trigger changes in target gene transcription. ZNF410, however, directly activates just one gene in human erythroid cells. This is supported by the following observations: 1) ZNF410 chromatin binding as measured by ChIP-seq is only seen at a total of eight regions, with by far the strongest signals occurring in the form of two peak clusters near the CHD4 gene. Failure to detect more ChIP-seq peaks was not a consequence of overlooking potentially bound regions because of mappability issues, such as those presented by repetitive elements, since inclusion of reads that map to multiple locations did not reveal additional binding sites. 2) Clusters of ZNF410 motifs such as those at the CHD4 locus are not found elsewhere in the genome. 3) At the non-CHD4 ZNF410-bound sites, signals were not only much weaker, but showed little or no signal in murine cells. Hence, ZNF410 chromatin occupancy is conserved only at the CHD4 locus. 4) Among the few ZNF410 bound genes, CHD4 was the only one whose expression was reduced upon ZNF410 loss or upon expression of dominant interfering ZNF410 constructs. 5) Forced expression of CHD4 almost completely restored γ-globin silencing and transcriptome in ZNF410 deficient cells. This also suggests that indirect, motif-independent binding to chromatin, which might escape detection by ChIP, would not have significant regulatory influence.

When interrogating data sets from 53 tissues, the ZNF410 and CHD4 mRNA levels are highly correlated, suggesting that ZNF410 may be generally limiting for CHD4 expression across tissues and cell lines. Notably, in luminal breast cancer cell lines, ZNF410 and CHD4 are the top co-essential genes, implying that they function in the same pathway (Depmap; https://depmap.org). Loss of ZNF410 does not completely abrogate CHD4 gene transcription. Consequently, the requirement of ZNF410 for CHD4 transcription is not absolute, implicating involvement of other factors in the regulation of the CHD4 gene. Whether ZNF410 has additional target genes in other tissues remains to be determined.

We are unaware of other transcriptional activators with single target genes, but there are cases of transcription factors with only very few target genes. For example, ZFP64 is an 11-zinc finger protein which binds most strongly to clusters of elements near the MLL gene, reminiscent of the ZNF410 motif clusters at the CHD4 locus (Lu et al., 2018). Yet ZFP64 displays thousands of additional high confidence ChIP-seq peaks even though it regulates only a small fraction of associated genes. The KRAB-ZFP protein Zfp568 is a transcriptional repressor that seems to only silence the expression of the Igf2 gene in embryonic and trophoblast stem cells, even though it occupies dozens of additional sites in the genome (Yang et al., 2017). Remarkably, deletion of the Igf2 gene in mice rescues the detrimental effects on gastrulation incurred upon Zfp568 loss, but embryonic lethality persists, implying the presence of additional Zfp568 repressed genes. Extraordinarily high gene selectivity has also been reported for transcriptional co-factors. For example, TRIM33, a cofactor for the myeloid transcription factor PU.1, has been shown to occupy only 31 genomic sites in murine B cell leukemia, and appears to preferentially associate with enhancers containing a high density of PU.1 binding sites (Wang et al., 2015). Transcription factors are normally employed at numerous genes, and spatio-temporal specificity is accomplished through combinatorial action with other transcription factors. However, the number of target genes for transcription factors and co-factors varies by three orders of magnitude (ENCODE Transcription Factor Targets). ZNF410 seems to have evolved to require motif clusters such as those found at the CHD4 locus to achieve such high levels of target gene specificity.

What accounts for the high selectivity of ZNF410 chromatin occupancy? 1) The human genome contains 434 perfect ZNF410 motif instances and 3677 similar ones if adding up all the motifs that are found under ZNF410 peaks, which is a much smaller number than that for the great majority of transcription factors (Srivastava and Mahony, 2020). Thus, motif scarcity is likely one determinant of target selectivity but obviously insufficient as the sole explanation. 2) ZNF410 binding site clusters are uniquely found at the CHD4 gene. If ZNF410 requires a cooperative mechanism for chromatin binding, this may explain lack of binding to the majority of single motifs. 3) The weak ZNF410 binding that is found at 6 sites containing a single motif is accompanied by the presence of active histone marks and signatures of open chromatin. It is possible that when exposed, single motifs might allow access to ZNF410 even if it is functionally inconsequential. Indeed, ZNF410 depletion or dominant interfering ZNF410 version elicited no transcriptional changes of the 6 ZNF410-bound genes with a single motif.

The ZF domain of ZNF410 is necessary and sufficient for DNA binding in vitro and in vivo. Crystallographic analysis of the ZF domain bound to DNA revealed a binding mode in which ZF1-ZF5 are contacting the consensus sequence in a 3’ to 5’ orientation with all five ZFs contacting DNA. EMSA experiments suggest, however, that four ZFs (either ZF1-4 or ZF2-5) are needed for efficient binding. Whether ZNF410 versions with fewer ZFs can achieve similar chromatin occupancy patterns as wild-type ZNF410 will be interesting to investigate in future experiments. Fluorescence polarization experiments measured the ZF domain-DNA interaction K_D_ at 22nM. Yet, this high affinity interaction appears insufficient to enable chromatin occupancy at virtually all single elements in the genome. Hence, the clustering of motifs may be required to convey efficient and high level chromatin binding. The molecular basis for binding co-operativity is unclear. Since the ZF domain displays no activation activity on its own and therefore might not interact with co-activator complexes, binding cooperativity might derive from the inherently synergistic effects of DNA binding domains when displacing histone-DNA interactions in nucleosomes (Adams and Workman, 1995; Oliviero and Struhl, 1991; Polach and Widom, 1996). Whether the spacing of the ZNF410 motifs allows for such synergistic behavior remains to be tested.

When overexpressed, the ZF domain acted in a dominant interfering manner by displacing endogenous ZNF410 from the CHD4 locus. The resulting reduction in CHD4 transcription was ∼65%-70%, comparable to that observed upon ZNF410 knockout. The expression of 6 other ZNF410 bound genes were unaffected, again illustrating ZNF410 specificity. One implication of this finding is that the transactivation function of ZNF410 resides outside the ZF domain, and that, by inference, the ZF domain may not be involved in co-activator recruitment. This contrasts with other zinc finger transcription factors, such as GATA1 where the ZF region can be multifunctional and not only bind DNA but also critical co-regulators (Campbell et al., 2013). Finally, according to our ChIP-seq experiments, the ZF domain binding profiles are very similar to full length ZNF410, suggesting that the ZNF410 chromatin binding specificity and affinity is determined solely by the ZF domain, and that other domains and associated cofactors contribute little if at all to ZNF410 binding.

Sequence variants at binding sites for the γ-globin repressors BCL11A and LRF (ZBTB7A) are linked to persistence of γ-globin expression into adulthood (Liu et al., 2018; Martyn et al., 2018). We interrogated GWAS Central databases as well as a sequencing database we generated (unpublished data) for SNPs within the CHD4 regulatory regions that might be linked to elevated HbF levels but found none. Given the large number of ZNF410 elements at the CHD4 locus, multiple elements would need to be lost in order to significantly affect CHD4 transcription. It is thus possible that motif clustering at the CHD4 locus provides robustness for the maintenance of CHD4 expression.

Complete CHD4 loss severely compromises hematopoiesis and erythroid cell growth (Sher et al., 2019; Xu et al., 2013; Yoshida et al., 2008). However, depletion of ZNF410 is well tolerated in erythroid cells and other hematopoietic lineages, which is likely due to the fact that CHD4 is not completely extinguished. This partial CHD4 reduction was sufficient to robustly de-repress the γ-globin genes, consistent with a prior study (Amaya et al., 2013). Notably, given the very limited global transcriptional changes upon ZNF410 depletion, this suggests that the γ-globin genes are especially sensitive to NuRD levels.

In sum, we identified ZNF410 as a highly specific regulator of CHD4 expression and γ-globin silencing. It might be possible to exploit this high transcriptional selectivity and target ZNF410 to raise fetal hemoglobin expression for the treatment of hemoglobinopathies.

## Data and Code Availability

The X-ray structures (coordinates and structure factor files) of ZNF410 ZF domain with bound DNA have been submitted to PDB under accession number 6WMI.

The RNA-seq and ChIP-seq data have been deposited to GEO database (GSE154963).

## Acknowledgments

The authors thank Dr. Shaun Mahony, Dr. Xing Zhang and members of the Blobel laboratory for helpful comments and discussions, and the CHOP flow cytometry core for help with cell sorting. The authors also thank the following core facilities and individuals at St. Jude Children’s Research Hospital: Flow Cytometry (Richard Ashmun, Jonathan Laxton and Stacie Woolard), Animal Resource Center (Chandra Savage), Center for advance genome editing (Shondra Miller and Shaina Porter), and Animal maintenance (Kalin Mayberry).

This work was supported by NIH grants R24DK106766 (G.A.B. and R.C.H.), R01HL119479 (G.A.B.), R35GM134744 (X.C.), CPRIT RR160029 (X.C.), Australian National Health and Medical Research Council APP1164920 (M.C.), a scholarship by Australian Government Research Training Program (L.C.L.), the St. Jude sponsored CRC consortium, and a fellowship by the Cooley’s anemia foundation (X.L.). We thank the Di Gaetano family for their generous support.

## Author Contributions

X.L. J.S., and G.A.B. conceived the study and designed the experiments. X.L., N.A., and O.A. performed experiments. J.D.G., and J.S. performed CRISPR-screen. R.R. performed protein purification, DNA binding assays and crystallization. R.R. and J.R.H. performed X-ray data collection and structure determination. X.C organized and designed the scope of the structural study. R.F. and T.M. performed mouse xenotransplantation experiments, and R.F., T.M. and M.J.W. analyzed the data. L.C.L and M.C carried out the band retardation experiments. X.L. Y.L., Z.Z., B.G., R.C.H. and K.Q. analyzed ChIP-seq and RNA-seq data. X.L. and G.A.B. analyzed data. X.L. and G.A.B. wrote the manuscript with input from all authors.

## Methods and materials

### Cell culture

HUDEP-2 cells were cultured and differentiated as described previously (Kurita et al., 2013). Briefly, StemSpan™ SFEM supplemented with 50ng/ml human SCF, 10μM dexamethasone, 1μg/ml doxycycline, 3IU/ml erythropoietin and 1% penicillin/streptomycin was utilized for routine cell maintenance. Cell density was kept at 0.1-1×10^6/ml. HUDEP-2 cells were differentiated for 6-7 days in IMDM supplemented with 50ng/ml human SCF, 3IU/ml erythropoietin, 2.5% fetal bovine serum, 250μg/ml holo-transferrin, 10ng/ml heparin, 10μg/ml insulin, 1μg/ml doxycycline and 1% penicillin/streptomycin.

Primary human CD34+ HSPCs from mobilized peripheral blood were purchased from the Fred Hutchinson Cancer Research Center. Human CD34+ HSPCs were differentiated using a three-phase culture system as described previously (Grevet et al., 2018). Briefly, IMDM supplemented with 3IU/ml erythropoietin, 2.5% human male AB serum, 10ng/ml heparin, and 10μg/ml insulin was used as base medium. For phase I medium, 100ng/ml human SCF, 5ng/ml IL-3, and 250μg/ml holo-transferrin were supplemented. For phase II medium, 100ng/ml human SCF and 250μg/ml holo-transferrin were supplemented. For phase III medium, 1.25mg/ml holo-transferrin was supplemented.

HEK293T cells were grown in DMEM supplemented with 10% fetal bovine serum, 2% penicillin/streptomycin, 1% L-glutamine and 100μM sodium pyruvate according to standard protocol.

K562 cells were cultured in IMDM supplemented with 10% fetal bovine serum and 1% penicillin/streptomycin.

G1E-ER4 cells are a sub-line of G1E cells, (derived from GATA1 KO murine embryonic stem cells (Weiss et al., 1997)), which express GATA1 fused to the ligand binding domain of the estrogen receptor (GATA1-ER) (Weiss et al., 1997). GATA1 activation and erythroid differentiation are induced by the addition of 100 nM estradiol to the media for 24 hours. G1E-ER4 were cultured in IMDM supplemented with 15% FBS, 1% penicillin/streptomycin, Kit ligand, monothioglycerol and epoetin alpha.

COS-7 cells were cultured in DMEM supplemented with 10% fetal bovine serum and 1% penicillin-streptomycin-glutamine (PSG). During passaging, adherent cells were dislodged after a 2-min incubation at 37 °C with PBS-EDTA (5 mM).

### Vector construction

SgRNAs were cloned into a lentiviral U6-sgRNA-EFS-GFP/mCherry expression vector (LRG, Addgene: #65656) by BsmBI digestion. The ZNF410 cDNA (clone ID: OHu10535), CHD4 cDNA (clone ID: OHu28780) were purchased from GenScript and were sub-cloned into a lentiviral vector pSDM101-IRES-GFP (from Dr. Patrick Grant lab). The ZF domain of ZNF410 was also sub-cloned into pSDM101-IRES-GFP vector. The N-terminal HA tag was introduced by PCR with primer. For EMSA assay, the ZNF410 full length or ZF domain was sub-cloned into mammalian expression vector pcDNA3.

### Lentiviral transduction

Lentivirus was produced as described previously (Grevet et al., 2018). Briefly, 10-20 ug of expression vectors, 5-10ug of pVSVG (pMD2.G) and 7.5-15 ug of psPAX2 package plasmids, and 80 ul of 1 mg/ml polyethylenimine (PEI) were mixed, incubated and added to the 10 cm plate HEK293T cells above 90% confluence, media was replaced 6-8 hr post transfection, virus was collected 24 hours and 48 hours post-transfection and pooled. For infection, virus-containing supernatant was mixed with the indicated cell lines with 8 ug/ml polybrene and 10 mM HEPES, and then spin at 2250 rpm for 1.5 hrs at room temperature. Infected HUDEP-2 or K562 cells were selected by mCherry+ or GFP+ cell sorting 48 hours post-infection. To control cDNA expression levels, the GFP+ low cells were sorted.

### RNP electroporation

Commercial sgRNAs were purchased from IDT or Synthego. To assemble the RNP complexes, 100 pmol sgRNA and 50 pmol SpCas9 protein (from IDT) were incubated at room temperature for 15 mins. CD34+ HSPCs (50k-100k) at Day 3-4 of phase I culture were electroporated using P3 Primary Cell 4D NucleofectorTM X Kit (from Lonza) with program DZ100 (Bak et al., 2018).

### RT-qPCR

RT-qPCR was performed as described previously (Grevet et al., 2018). Briefly, total RNAs were purified using the RNeasy Plus Mini Kit (Qiagen) including an on-column DNAse treatment using RNase-free DNase set (Qiagen) to remove genomic DNA. Reverse transcription was accomplished using iScript Supermix (Bio-Rad). qPCR reactions were prepared with Power SYBR Green (ThermoFisher Scientific). Quantification was performed using the ΔΔC_T_ method. Primers used for RT-qPCR are listed in Table S1.

### COS cell transfections and nuclear extractions

Nuclear extracts were prepared from COS-7 cells transiently overexpressed with ZNF410 full-length and ZNF410 ZF1-5 plasmids. Fugene 6 (Promega) was used to transfect 5 μg of vector into 100 mm plates of COS-7 cells. A pcDNA3 empty vector was transfected as a control. Cells were harvested 48 h after transfection and nuclear extracts prepared as previously described (Andrews and Faller, 1991). The mammalian expression vectors used are presented in Table S3.

### In vivo transplantation of CD34+ HSPCs

Xenotransplantation experiments were carried out as previously described (Metais et al., 2019). Briefly, ZNF410 edited or control CD34+ HSPCs were administered at a dose of 0.4 million per NBSGW mouse (The Jackson Laboratory) by tail-vain injection at aged 8-12 weeks. Chimerism post-transplantation was assessed by flow analysis at 8 weeks in the periphery and at 16 weeks in the bone marrow at the time of euthanasia. Cell linage composition was determined in the bone marrow using human-specific antibodies, and different lineages were sorted by a FACSAria III cell sorter. CD34+ HSPCs were isolated with magnetic beads using the human-specific CD34 MicroBead Kit UltraPure, human (Miltenyi Biotec Inc). See table S4 for antibodies used in flow cytometry.

### Indel analysis

Next-generation sequencing (NGS) was used for indel analysis as previously described (Metais et al., 2019). Briefly, NGS libraries were prepared with a 2-step PCR protocol. In the first step, the targeted genomic sites were amplified by PCR with Phusion Hot Start Flew 2x Master Mix (New England BioLabs) and primers with partial Illumina sequencing adaptors. In the second step, PCR was performed with a KAPA HiFi HotStart ReadyMix PCR Kit (Roche) to add Illumina sequencing adapters (P5-dual-index and P7-dual-index) to the purified PCR product from the first step. The Illumina MiSeq platform was used to generate FASTQ sequences with 150 bp paired-end reads, and these reads were analyzed by joining paired reads and analyzing amplicons, using CRISPResso for indel measurement. See table S5 for NGS sequencing primers.

### EMSAs

EMSAs were carried out as previous described (Crossley et al., 1996). Oligonucleotides used in the synthesis of radiolabelled probes are listed in Table S6. The sense oligonucleotide was labelled with [γ-^32^P]-adenosine triphosphate (Perkin Elmer) and boiled at 100 °C for 1 min before addition of the antisense oligonucleotide and annealing of probe via slow cooling from 100 °C to room temperature. Probes were purified using Quick Spin Columns for Radiolabelled DNA Purification (Roche). Nuclear extracts were harvested from COS-7 cells and samples were loaded on a 6% native polyacrylamide gel in TBE buffer (45 mM Tris, 45 mM boric acid, 1 mM EDTA). A ‘COS empty’ control lane was included to show binding of any background endogenous protein to the probe. Recognition and super-shifting of FLAG-ZNF410 overexpression constructs was achieved by using the anti-FLAG monoclonal antibody (Sigma). Gels were run at 250 V for 1h 45 min at 4 °C then dried under vacuum. Gels were exposed overnight with a FUJIFILM BAS CASETTE2 2025 phosphor screen and imaged using the Typhoon™ FLA 9500 Laser Scanner.

### HbF staining and flow cytometry

HbF staining and flow analysis were performed as described previously (Grevet et al., 2018).

### CRISPR sgRNA library generation and screen

SgRNA library targeting human transcription factors was described previously (Lu et al., 2018). CRISPR-Cas9 screen was performed as described previously (Grevet et al., 2018).

### Western Blot

Western blotting analysis was carried out according to standard protocol with the following antibodies: ZNF410 (1:500, Proteintech, Cat. # 14529-1-AP), CHD4 (1:500, Proteintech, Cat. # 14173-1-AP), CHD4 (1:500, Cell Signaling Cat. # 11912S), GATAD2A (1;1000, Bethyl Laboratories, Cat. # A302-358A-T), HDAC2 (1;1000, Bethyl Laboratories, Cat. # A300-705A-T), MBD2 (1;1000, Bethyl Laboratories, Cat. # A301-633A-T), MTA2 (1;1000, Bethyl Laboratories, Cat. # A300-395A-T), HA (1:1000, Cell Signaling Cat. #3724), FLAG (1:1000, Sigma, Cat. # F1804), BCL11A (1:1000, Abcam, Cat. #19487), LRF (1:1000, eBioscience Cat. #13E9), GATA1 (1:1000, Santa Cruz, Cat. #sc-265), γ-globin (1:1000, Santa Cruz, Cat. #sc-21756), β-actin (1:1000, Santa-Cruz, Cat. #sc-47778). Secondary antibodies: anti-rabbit (1:10,000, GE Healthcare, Cat. #NA934V); anti-mouse (1:10,000, GE Healthcare, Cat. #NA931V) anti-Rat (1: 5,000, ThermoFisher Scientific, Cat. #31470), anti-Hamster (1: 5,000, ThermoFisher Scientific, **Cat. #** PA1-32045).

### RNA-Seq

Total RNAs were purified as described above. Sequencing libraries were then constructed using 100 ng of purified total RNA using the ScriptSeq Complete Kit (Illumina cat# BHMR1224) according to manufacturer’s protocol. In brief, the RNA was subjected to rRNA depletion using the Ribo-Zero removal reagents and fragmented. First strand cDNA was synthesized using a 5’ tagged random hexamer, and reversely transcribed, followed by annealing of a 5’ tagged, 3’-end blocked terminal-tagged oligo for second strand synthesis. The Di-tagged cDNA fragments were purified, barcoded, and PCR-amplified for 15 cycles.

The size and quality of each library were then evaluated by Bioanalyzer 2100 (Agilent Techniologies, Santa Clara, CA), and quantified using qPCR. Libraries were sequenced in paired-end mode on a NextSeq 500 instrument to generate 2 × 76 bp reads using Illumina-supplied kits. The sequence reads were processed using the ENCODE3 long RNA-seq pipeline (https://www.encodeproject.org/pipelines/ENCPL002LPE/). In brief, reads were mapped to the human genome (hg38 assembly) using STAR, followed by RSEM for gene quantifications.

### RNA-Seq data analysis

The normalized FPKM (fragments per kilo base per million mapped reads) for each gene was averaged in 2 replicates and then filtered to keep those with average FPKM at least 10 in both HUDEP-2 cells and primary erythroblasts, resulting in ∼5000 high abundant genes each cell type for further analysis. Log2 fold-change was calculated from FPKM of sgRNA targeting ZNF410 compared to control sgRNA (non-targeting sgRNA) using the DESeq2 method, and top changed genes were selected with fold-change at least 1.5 and p-value<0.05. Commonly changed genes in both independent sgRNAs were considered to be significant. Scatter plots were generated using ggplot2 in RStudio for all expressed genes (FPKM>5).

### ChIP-seq

HUDEP-2 cells at Day 3 differentiation, primary human CD34+ cells at Day 9 differentiation and G1E-ER4 cells at 24 hours differentiation were crosslinked with 1% formaldehyde at room temperature for 10 min and quenched by the addition of glycine. ChIP experiments were performed as previously described (Hsu et al., 2017). ZNF410 (Proteintech, Cat. # 14529-1-AP), HA (Sigma, Cat. # 11815016001) and H3K27ac (Abcam, Cat. # ab4729) antibodies were used for ChIP. ChIP-seq libraries were prepared using TruSeq ChIP-seq Sample preparation Kit (part# IP-202-1012) according to the manufacturer’s instructions. Reads were aligned with Bowtie2 local alignment to allow the mapping of indels (Langmead and Salzberg, 2012). All ChIP-seq experiments were performed in two biological replicates. ChIP-qPCR was performed with Power SYBR Green (ThermoFisher). Primers used for ChIP-qPCR are listed in Table S7.

### ZNF410 ChIP-Peak calling and de novo motif analysis

Reads were aligned against reference genome hg38 using Bowtie2 (v2.2.9) and the default parameters. Alignments with MAPQ score lower than 10 and PCR duplicates were removed using Samtools (v0.1.19). Reads aligned to mitochondria, random contigs and ENCODE blacklisted regions were also removed for downstream analysis. Genome coverage files were generated and normalized to 1 million reads per library using bedtools (v2.25.0), and then converted to bigwig format for visualization using the UCSC Toolkit. Peaks were called using MACS2 (v2.1.0) and a 0.05 q-value cutoff. The final peaks were those overlapped by both ZNF410 replicates but not in control replicates (empty vector and knock-out samples), then manually filtered to exclude peaks near centromere/telomere regions that did not look like peaks on genome browser (total number reduced from 38 to 8). The final peaks were extended by 1kb on both ends for de novo motif analysis using the HOMER tool, and the top hit motif was scanned across the entire genome using HOMER. We also scanned the human and mouse genome for motif pattern of CATCCCATAATA and other similar motifs using EMBOSS fuzznuc (v6.5.7.0). Read density plot and heatmap around selected peaks were generated using Deeptools (version 2.5.7, “computeMatrix” and “plotHeatmap”).

### HPLC

∼1 million cells were lysed in water for 10 mins, vertex 10 sec every 5 mins at RT. Hemolysates were then cleared by centrifugation at 15,000 rpm, 10 mins and analyzed for identity and levels of hemoglobin variants (HbF and HbA) by cation-exchange high-performance liquid chromatography (HPLC). Hitachi D-7000 Series (Hitachi Instruments, Inc., San Jose, CA), and weak cation-exchange column (Poly CAT A: 35 mm x 4.6 mm, Poly LC, Inc., Columbia, MD) were used. Hemoglobin isotype peaks were eluted with a linear gradient of phase B from 0% to 80% at *A*_410nm_ (Mobile Phase A: 20 mM Bis-Tris, 2 mM KCN, pH 6.95; Phase B:20 mM Bis-Tris, 2 mM KCN, 0.2 M sodium chloride, pH 6.55). Cleared lysates from normal human cord blood samples (high HbF content), as well as a commercial standard containing approximately equal amounts of HbF, A, S and C (Helena Laboratories, Beaumont, TX), were utilized as reference isotypes.

### Wright-Giemsa staining

∼100,000 cells were spun onto glass slides with Cytospin4 (ThermoFisher Scientific) at 1,200rpm for 3 min. Slides were allowed to dry for 5 minutes at RT, followed by staining with May Grünwald (Sigma Aldrich) for 2 minutes and then by 1:20 diluted Giemsa stain (Sigma Aldrich) for 10 minutes. The stained slides were rinsed twice in water and then allowed to dry for 10 minutes before a coverslip was sealed on the preparation with Cytoseal 60 (Thermo Scientific). The images were captured with Olympus BX60 microscope at 10X resolution using Infinity software (Lumenera corporation).

### Protein expression and purification

The fragment of Human ZNF410 (NP_001229855.1) comprising of five zinc finger domains ZF1-5 (residues 217-366) was cloned into pGEX-6P-1 vector with a GST fusion tag (pXC2180). The plasmid was transformed into *Escherichia coli* strain BL21-Codon-plus(DE3)-RIL (Stratagene). Bacteria was grown in LB broth in a shaker at 37°C until reaching the log phase (A_600nm_ between 0.4 and 0.5), the shaker temperature was then set to 16°C and 25 μM ZnCl_2_ was added to the cell culture. When the shaker temperature reached 16°C and A_600nm_ reached *∼*0.8, the protein expression was induced by the addition of 0.2 mM isopropyl-β-D-thiogalactopyranoside with subsequent growth for 20 h at 16°C. Cell harvesting and protein purification were carried out at 4°C through a three-column chromatography protocol (Patel et al., 2016a), conducted in a BIO-RAD NGC™ system. Cells were collected by centrifugation and pellet was suspended in the lysis buffer consisting of 20 mM Tris-HCl, pH 7.5, 500 mM NaCl, 5% glycerol, 0.5 mM tris(2-carboxyethl)phosphine (TCEP) and 25 μM ZnCl_2_. Cells were lysed by sonication and 0.3% (w/v) polyethylenimine was slowly titrated into the cell lysate before centrifugation (Patel et al., 2016a). Cell debris was removed by centrifugation for 30 min at 47,000 xg and the supernatant was loaded onto a 5 ml GSTrap column (GE Healthcare). The resin was washed by the lysis buffer and bound protein was eluted with elution buffer of 100 mM Tris-HCl, pH 8.0, 500 mM NaCl, 5% glycerol, 0.5 mM TCEP and 20 mM reduced form glutathione. The GST fusion were digested with PreScission protease (produced in-house) to remove the GST fusion tag. The cleaved protein was loaded onto a 5 ml Heparin column (GE Healthcare). The protein was eluted by a NaCl gradient from 0.25 to 1 M in 20 mM Tris-HCl, pH 7.5, 5% glycerol and 0.5 mM TCEP. The peak fractions were pooled, concentrated and loaded onto a HiLoad 16/60 Superdex S200 column (GE Healthcare) equilibrated with 20 mM Tris-HCl, pH 7.5, 250 mM NaCl, 5% glycerol and 0.5 mM TCEP. The protein was frozen and stored at -80°C.

### DNA binding assays

Fluorescence polarization (FP) method was used to measure the binding affinity using a Synergy 4 Microplate Reader (BioTek). Aliquots (5 nM) of 6-carboxy-fluorescein (FAM)-labeled DNA duplex (FAM-5’-CACA TCC CAT AAT AATG-3’ and 3’-GTGT AGG GTA TTA TTAC-5’) and control (FAM-5’-TCC ACT GCC AGG ACC TTT-3’ and 3’-GGT GAC GGT CCT GGA AAA-5’) was incubated with varied amount of proteins (0 to 2.5 μM) in 20 mM Tris-HCl, pH 7.5, 300 mM NaCl, 5% glycerol and 0.5 mM TCEP for 10 min at room temperature. The data were processed using Graphpad Prism (version 8.0) with equation [mP] = [maximum mP] × [C] / (KD +[C]) + [baseline mP], in which mP is millipolarization and [C] is protein concentration. The K_D_ value for each protein–DNA interaction was derived from two replicated experiments.

Electrophoretic mobility shift assay (EMSA) was performed with the same set of samples used in the FP assay for 10min at room temperature. Aliquots of 10 μl of reactions were loaded onto an 8% native 1x TBE polyacrylamide gel and run at 150V for 20 min in 0.5x TBE buffer. The gel was imaged using a ChemiDoc Imaging System (BIO-RAD).

### Crystallography

The ZF-DNA complex was prepared by mixing 0.9 mM ZF1-5 fragment and double-stranded DNA oligo (annealed in buffer containing 10 mM Tris-HCl, pH 7.5, and 50 mM NaCl) with molar ratio 1:1.2 of protein to DNA on ice for 30 min incubation. The protein-DNA complex crystals were grown using the sitting drop vapor diffusion method via an Art Robbins Gryphon Crystallization Robot at 19 °C with a well solution of 0.2 M ammonium formate and 20% polyethylene glycol 3350. Crystals were flash frozen using 20% (v/v) ethylene glycol as the cryo-protectant. The X-ray diffraction data were collected at SER-CAT 22-ID beamline of the Advanced Photon Source at Argonne National Laboratory utilizing a X-ray beam at 1.0 Å wavelength and processed by HKL2000 keeping Friedel mates separate (Otwinowski et al., 2003).

The resultant dataset for ab initio phasing was examined using the PHENIX Xtriage module (Adams et al., 2002) which reported a very good anomalous signal to 5.6 Å. The PHENIX AutoSol module (Terwilliger et al., 2009) identified the space group being *P*6_2_ and found all 10 zinc atom positions (5 per each of two molecules in asymmetric unit) with a Figure-Of-Merit of 0.48 and gave a density modified map with an R-factor of 0.34 at 5 Å data. Insertion of these zinc positions into AutoSol and utilizing the full resolution of the dataset gave a Figure-Of-Merit of 0.28 and a density modified map with an R-factor of 0.34. DNA duplex and zinc fingers bound in the major groove could easily be identified for the resultant map. The AutoBuild module of PHENIX was utilized for model building, and manual fitting of the protein and the DNA duplex was completed with COOT (Emsley and Cowtan, 2004), which was also utilized for corrections between PHENIX refinement rounds. Structure quality was analyzed during PHENIX refinements and finally validated by the PDB validation server. Molecular graphics were generated by using PyMol (Schrödinger, LLC).

## Supplemental Figures

**Figure S1.**
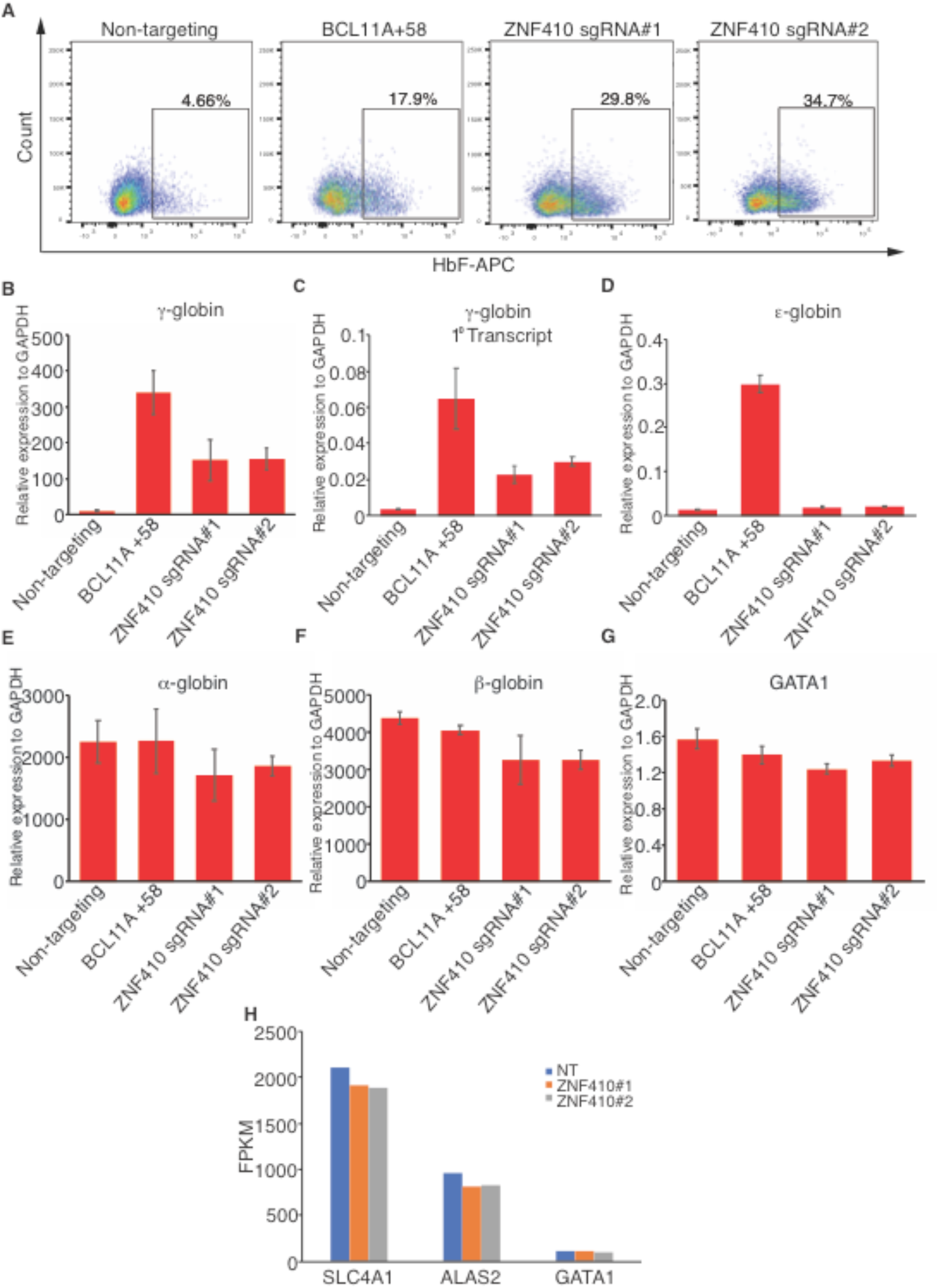
HbF flow cytometric analysis and RT-qPCR in HUDEP-2 cells, related to Figure 1. (A)Representative flow cytometric analysis of differentiated HUDEP-2 cells stained with anti-HbF antibody. Positive control: sgRNA against BCL11A +58: Negative control: Non-targeting sgRNA. (B-G) mRNA levels of γ-globin, γ-globin primary transcripts, ε-globin, α-globin, β-globin and GATA1 by RT-qPCR. Results are shown as mean ± SD (n=3). GAPDH was used for normalization. 1^0^ transcript: primary transcript. (H)Expression levels of SLC4A1(Band3), ALAS2 and GATA1 in differentiated HUDEP-2 cells transduced with indicated sgRNAs by RNA-seq analysis. NT: non-targeting.

**Figure S2.**
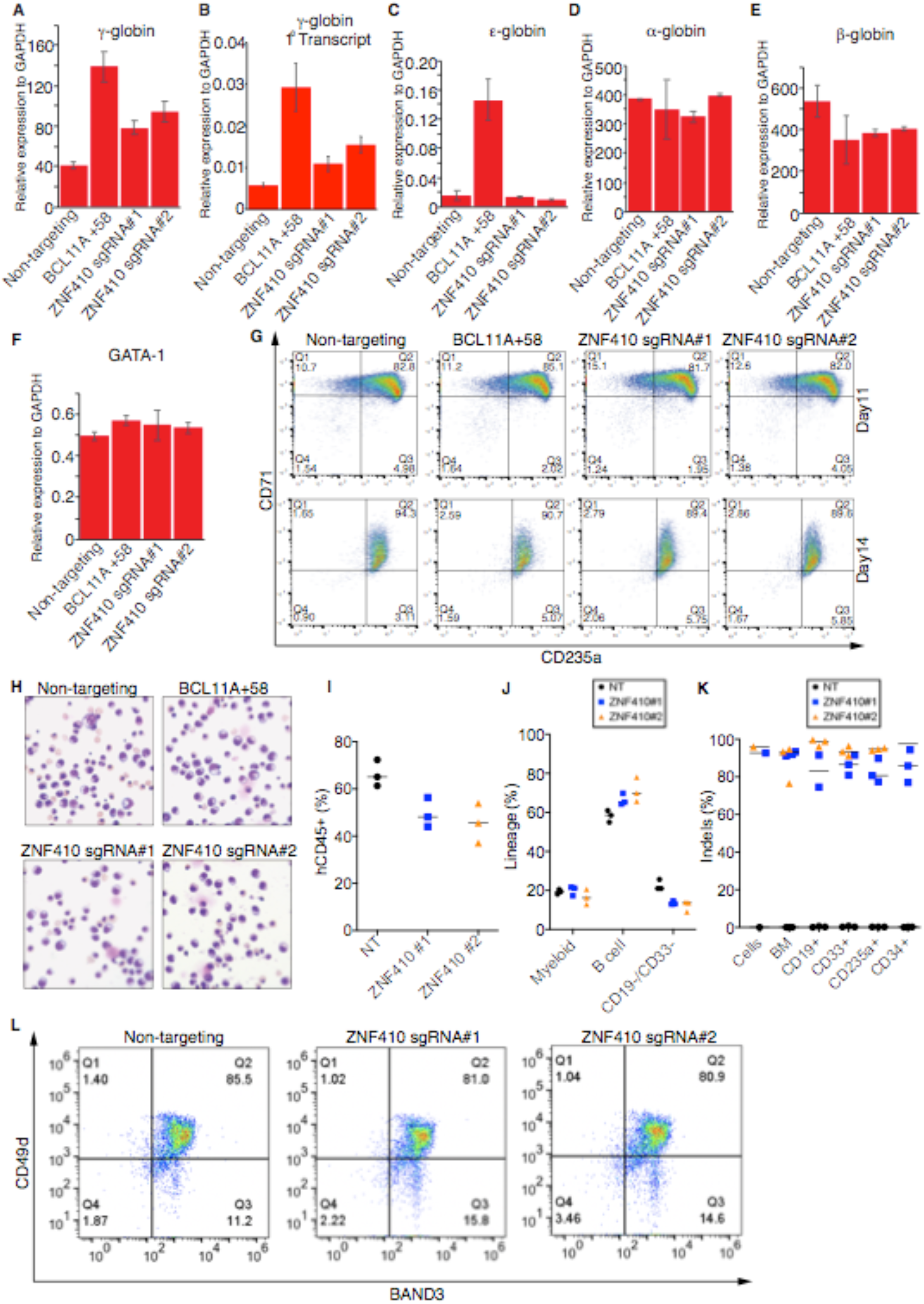
ZNF410 depletion induces HbF with minimal impact on erythroid maturation in vitro and in vivo, related to Figure 2. (A-F) mRNA levels of γ-globin, γ-globin primary transcripts, ε-globin, α-globin, β-globin and GATA1 by RT-qPCR in cultured primary human erythroblasts on day 12 of differentiation. Positive control: sgRNA against BCL11A +58: Negative control: Non-targeting sgRNA. Results are shown as mean ± SD (n=3). GAPDH was used for normalization. (G) Representative flow cytometric analysis of erythroid maturation markers CD71 and CD235a in cultured primary human erythroblasts on day 11 and day 14 of differentiation. (H) Wright-Giemsa staining in cultured primary human erythroblasts at day 16 of differentiation. (I) Normalized human chimerism in bone marrow from NBSGW mice at 16 weeks after transplantation, shown as percentage of human (h) CD45+ cells. (J) Human myeloid (CD33+), B cells (CD19+) and other cell types (CD19-/CD33-) shown as percentages of the human CD45+ population in bone marrow (BM) from NBSGW mice at 16 weeks after transplantation. (K) Indels measured by next generation sequencing (NGS) in input cells, bone marrows (BM) and specific hematopoietic lineages including B cells (CD19+), myeloid (CD33+), erythroblasts (CD235a+) and hematopoietic stem cells (CD34+). (L) Representative flow cytometric analysis of erythroid maturation markers CD49d/Band3 in human CD235a+ erythroblasts from recipient mouse bone marrow. For (I-K) panels, each dot in graphs represents a separate mouse. n=3 mice per sgRNA. NT: non-targeting.

**Figure S3.**
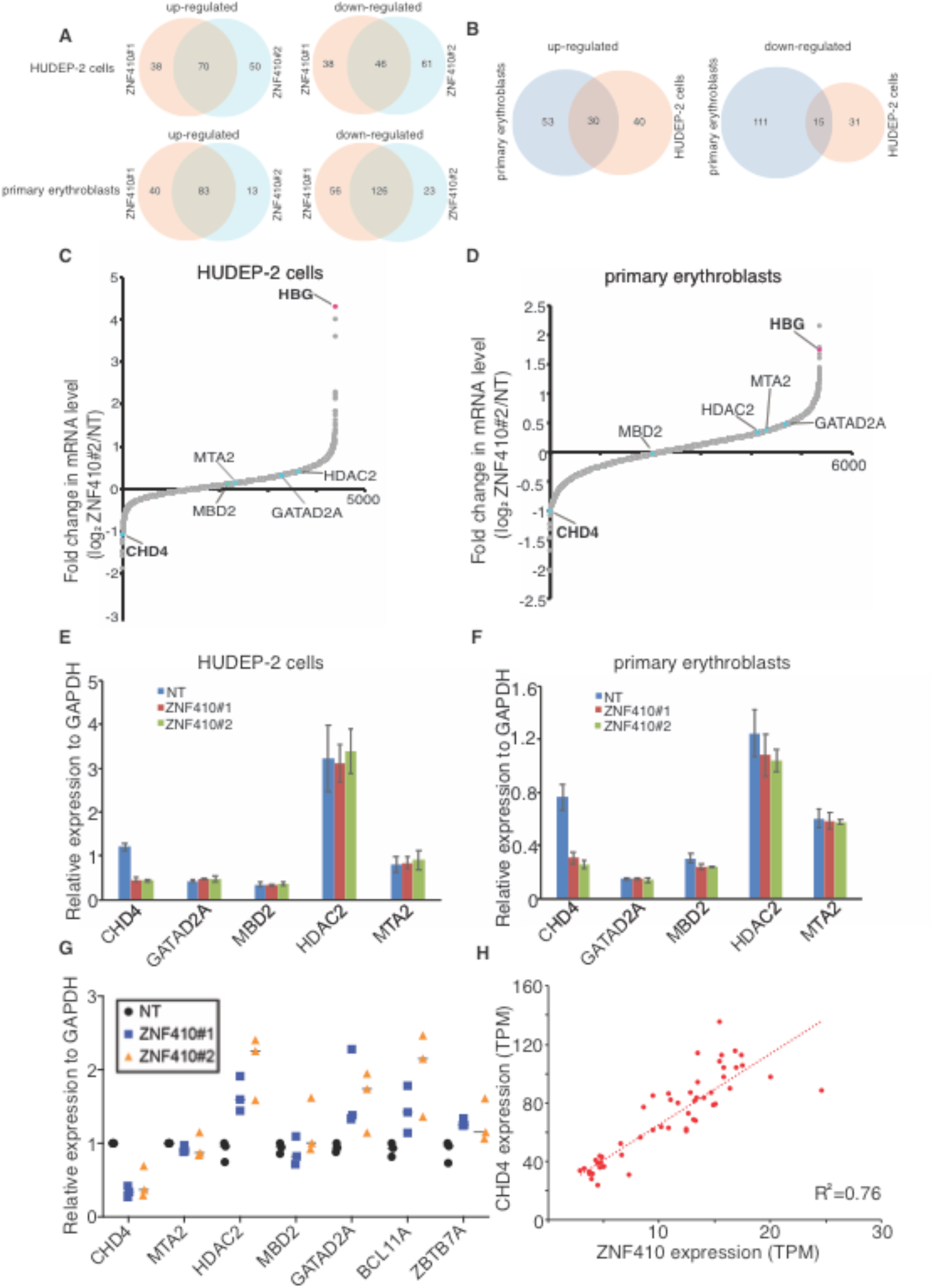
ZNF410 depletion diminishes CHD4 transcription, related to Figure 3. (A) Venn diagrams of the commonly upregulated and downregulated genes in both ZNF410 sgRNAs in HUDEP-2 cells or primary erythroblasts. (B) Venn diagrams of the commonly upregulated and downregulated genes in both HUDEP-2 and primary erythroblasts. (C-D) RNA-seq analysis of HUDEP-2 cells transduced with ZNF410 sgRNA#2 and primary erythroblasts with ZNF410 sgRNA#2. Plotted is the average fold-change in mRNA levels of two biological replicates. FPKM value was used to calculate fold change for each gene. NuRD complex subunits and γ-globin genes are indicated. NT: non-targeting. (E-F) mRNA levels of the NuRD complex subunits including CHD4, HDAC2, GATAD2A, MBD2 and MTA2 by RT-qPCR in HUDEP-2 cells and primary erythroblasts. Results are shown as mean ± SD (n=2). GAPDH was used for normalization. NT: non-targeting. (G) mRNA levels of the NuRD complex subunits and BCL11A and ZBTB7A by RT-qPCR in human CD235a+ erythroblasts from recipient NBSGW mouse bone marrow. Each dot in graph represents a separate mouse. n=3 mice per sgRNA. GAPDH was used for normalization. NT: non-targeting. (H) Correlation of ZNF410 and CHD4 mRNA levels across 53 human tissues. TPM: transcripts per kilobase million. Each dot indicates one tissue. Expression data were obtained from the Genotype-Tissue Expression database.

**Figure S4.**
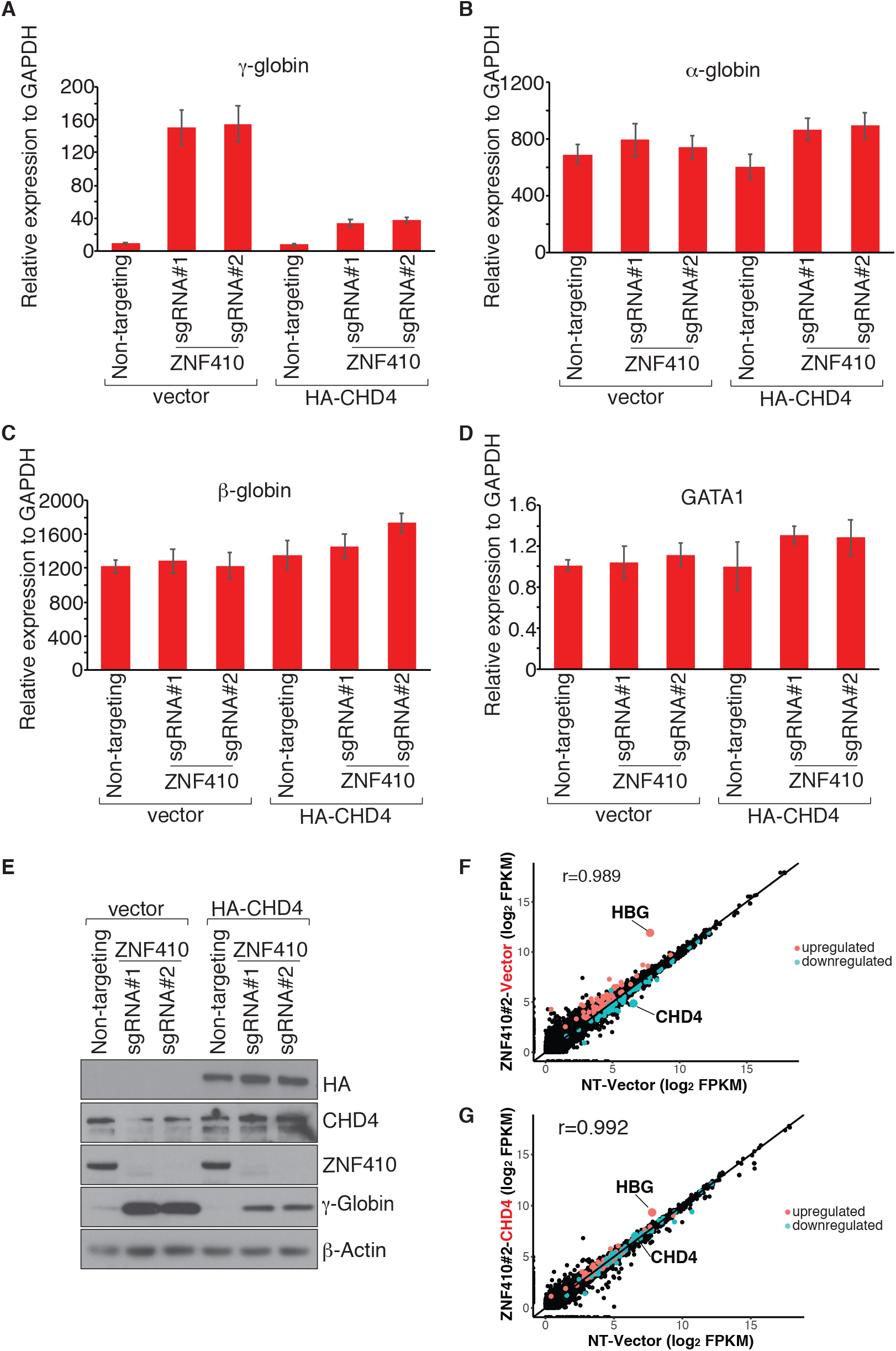
Re-introduction of CHD4 cDNA restores the silencing of HbF and transcriptome in ZNF410 deficient HUDEP-2 cells, related to Figure 3. (A-D) mRNA levels of γ-globin, α-globin, β-globin and GATA1 by RT-qPCR in ZNF410 deficient HUDEP-2 cells transduced with lentiviral vector containing CHD4 cDNA or empty vector. Results are shown as mean ± SD (n=2). GAPDH was used for normalization. (E) Western blot using whole-cell lysates from ZNF410 deficient HUDEP-2 cells with empty vector or forced CHD4 expression. HA-CHD4: N-terminal HA tagged CHD4. (F) Scatter plot of RNA-seq analysis in ZNF410 deficient HUDEP-2 cells (by ZNF410 sgRNA#2) with empty vector. Each gene is depicted according to averaged FPKM value from 2 biological replicates. r: Pearson’s correlation coefficient. NT: non-targeting. (G) Scatter plot of RNA-seq analysis in ZNF410 deficient HUDEP-2 cells (by ZNF410 sgRNA#2) with re-introduction of CHD4 cDNA. Each gene is depicted according to averaged FPKM value from 2 biological replicates. r: Pearson’s correlation coefficient. NT: non-targeting.

**Figure S5.**
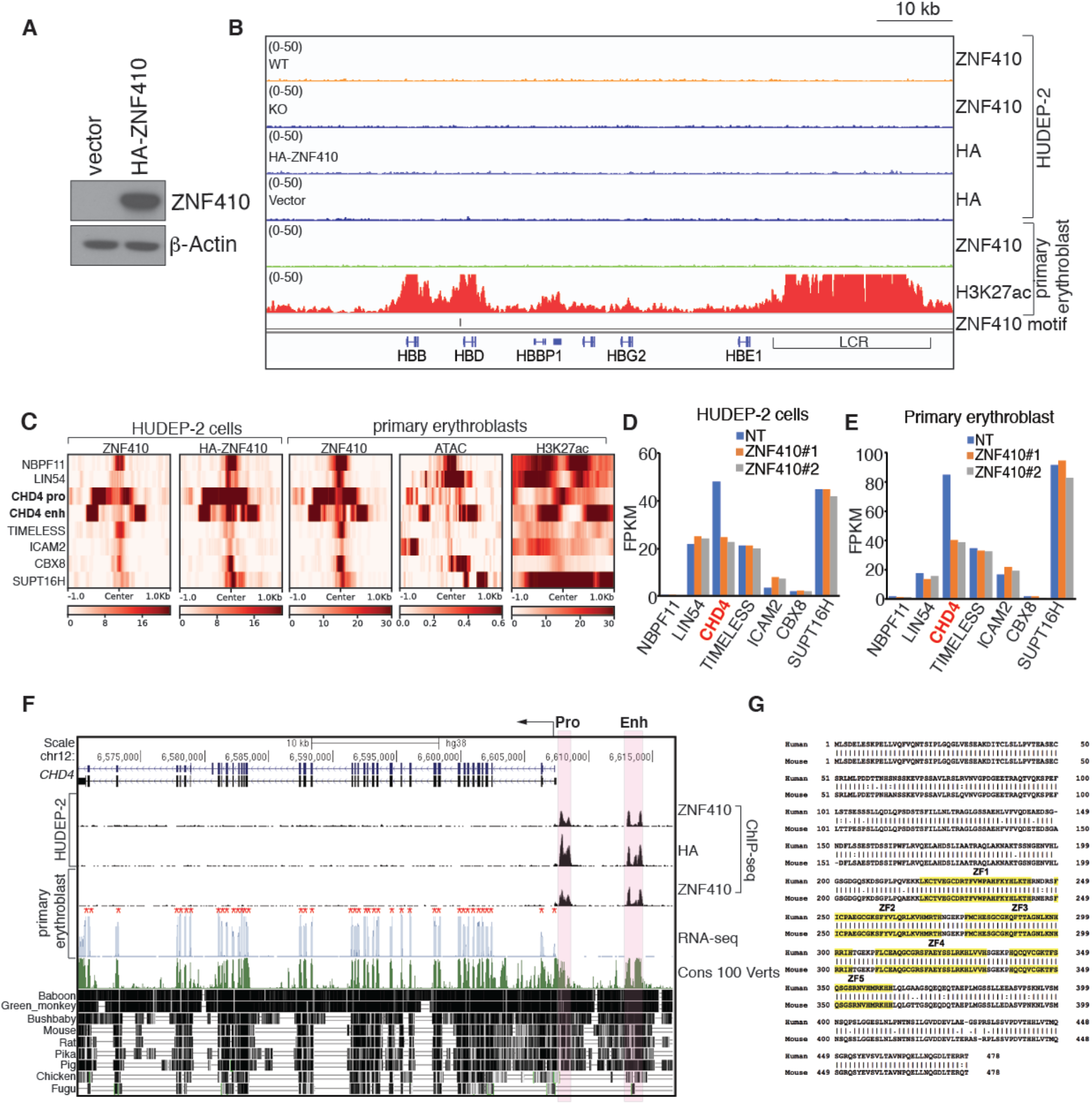
ZNF410 binds to the CHD4 locus in an evolutionarily conserved manner, related Figure 4. (A) Immunoblot analysis confirming overexpression of HA-ZNF410 in HUDEP-2 cells. (B) Gene track of endogenous ZNF410, HA-ZNF410 and H3K27ac ChIP-seq occupancy at the β-globin locus. The enhancer (LCR) is highlighted with line. LCR: local control region. (C) Heatmaps of the signal intensities of the 8 ZNF410 bound regions from endogenous ZNF410, HA-ZNF410, H3K27ac ChIP-seq and ATAC-seq in HUDEP-2 or primary erythroblasts. ATAC-seq were obtained from published data (Ludwig et al., 2019). (D-E) Expression levels of the 7 ZNF410 bound genes in HUDEP-2 cells transduced with indicated sgRNAs (F) and primary erythroblasts with indicated sgRNAs (G) by RNA-seq analysis. NT: non-targeting. (F) PhastCons (from 0 to 1) estimates of evolutionary conservation among 100 vertebrates at the CHD4 gene locus. The CHD4 promoter and enhancer are shaded in orange. CHD4 exons are marked by red * in RNA-seq track. ZNF410 ChIP-seq tracks show ZNF410 binding at the CHD4 promoter and enhancer. Pro: promoter, Enh: enhancer. Cons: conservation, Verts: vertebrates. (G) Alignment of human and mouse ZNF410 protein sequence. Identical residues are linked by vertical line. ZF (zinc finger) residues are shaded in yellow.

**Figure S6.**
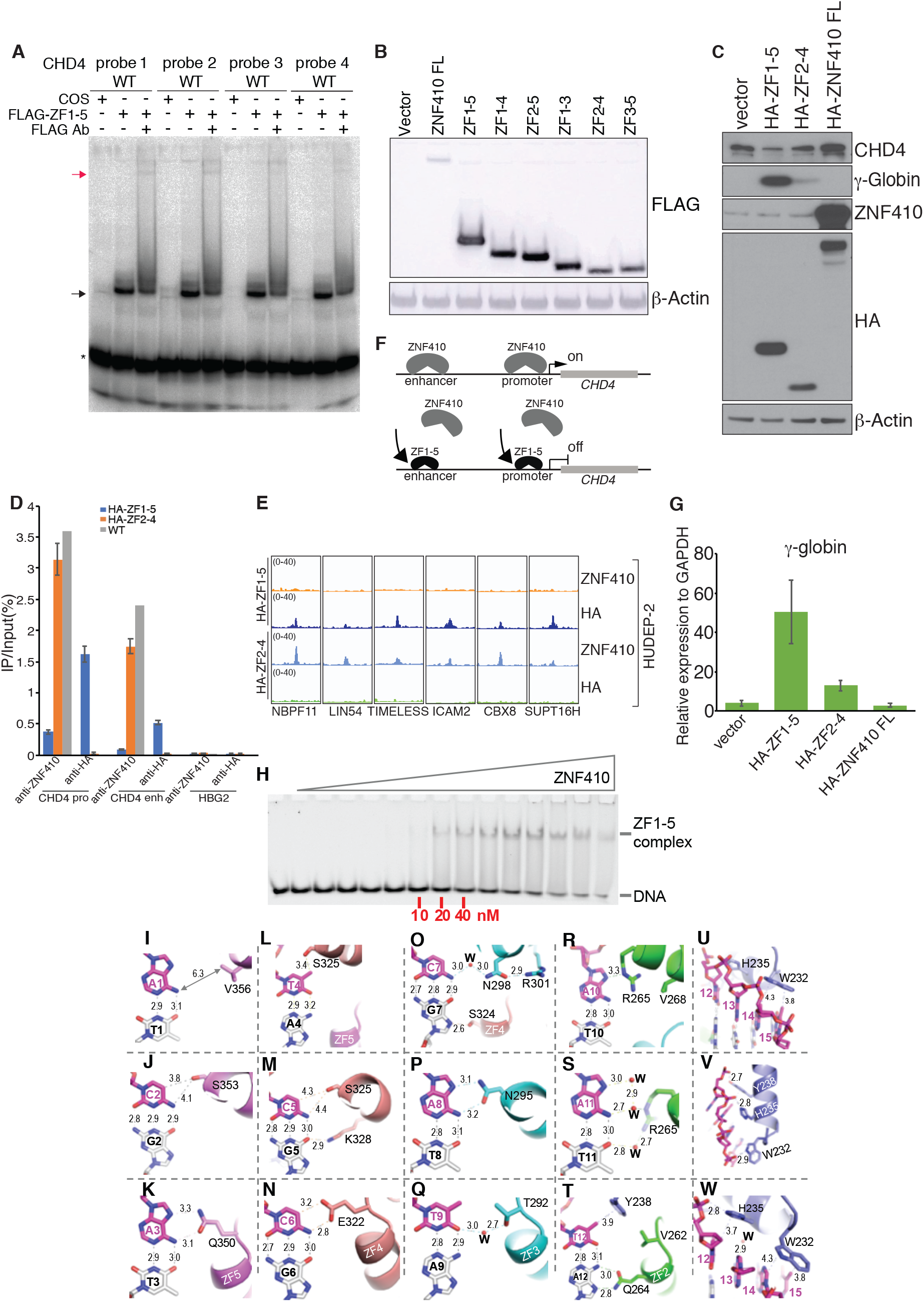
The ZF domain of ZNF410 is sufficient for DNA binding in vitro and in vivo, related to Figure 5 and Figure 6. (A) The ZF domain of ZNF410 binds to the four motifs from the CHD4 promoter and enhancer sites. Ab, antibody. Black arrow: ZF domain-probe complex; red arrow: FLAG antibody-ZF domain-probe complex. * : free probes. (B) Western blot analysis of FLAG-ZNF410 constructs expressed in COS-7 cells. Control: empty vector. All constructs were N-terminal FLAG tagged. (C) Western blot analysis using whole-cell lysates from differentiated HUDEP-2 cells overexpressing HA-ZF1-5, HA-ZF2-4 or HA-ZNF410 FL. (D) ZNF410 and HA ChIP-qPCR in WT HUDEP-2 cells, HA-ZF1-5 or HA-ZF2-4 overexpressed HUDEP-2 cells. HBG2 region serves as negative control. Results are shown as mean ± SD (n=2 for HA-ZF1-5 and HA-2-4 overexpressed cells) (E) ChIP-seq tracks of endogenous ZNF410, HA-ZNF410 at the 6 ZNF410 bound regions in HUDEP-2 cells overexpressing indicated constructs. (F) Diagram of HA-ZF1-5 displacing endogenous ZNF410 binding. (G) γ-globin levels measured by RT-qPCR in differentiated HUDEP-2 cells overexpressing indicated constructs. Empty vector serves as control. Results are shown as mean ± SD (n=2). GAPDH was used for normalization. (H) EMSA of ZNF410 ZF1-5. The maximum protein concentration used were 0.5 μM (the right most lane) followed by 2-fold serial dilutions. (I-K)ZF5 interactions with base pairs 1-3. V356 is too far from A1 to make direct contact. S353 makes van der Waals contacts with C2. Q350 interacts with A3. (L-N) ZF4 interactions with base pairs 4-6. S325 forms a weak hydrogen bond with T4, and makes van der Waals contact with C5. E322 interacts with C6. (O-Q) ZF3 interactions with base pairs 7-9. N298 conducts a water mediated interaction with C7. S324 interacts with G7. N295 interacts with A8. T292 conducts a water mediated interaction with T9. (R-T) ZF2 DNA interaction with base pairs 10-12. R265 forms a weak hydrogen bond with A10 and mediates a water network connecting with A:T base pair at position 11. Q264 interacts with A12. Y238 of ZF1 forms a π–methyl interaction with T12. (U-W) ZF1 interactions with base pair 12-15. W232 forms van der Waals contact with A14 and T15. H235, Y238 and W232 form polar interactions with DNA backbone phosphate groups.

## Supplementary Text

ZF2 and ZF3 have water-mediated contacts via R265 of ZF2 and T292 and N298 of ZF3 (Figures S6O, S6Q and S6S). We note that Arg is the most common mechanism for Gua recognition and Asn for Ade recognition (as shown above) (Luscombe et al., 2001; Patel et al., 2016b; Patel et al., 2018). When N298 of ZF3 meets C:G at position 7 and R265 of ZF2 meets A:T at position 11, both side chains of N298 and R265 moves away from DNA bases and allow water molecules to diffuse in. N298 forms a H-bond with R301 (Figure S6O), whereas R265 rotates its side-chain guanidine group bridging between two neighboring A10 and A11 (Figures S6R and S6S). Similarly, histidine is the next favorable residue for Gua recognition, when H235 of ZF1 meets an A:T base pair at position 14, H235 simply rotates its side chain imidazole ring and forms charge-charge interaction with a backbone phosphate group (Figure S6V). The smaller serine residues of ZF4 and ZF5 have non-specific contacts with C2 (via S353 of ZF5), T4 and C5 (via S325 of ZF4), and G7 (via S324 of ZF4) (Figures S6J, S6L, S6M and S6O), because serine can act as both an H-bond donor and acceptor and might accommodate alternative bases (Patel et al., 2017).

## Supplemental tables

**Table S1.**
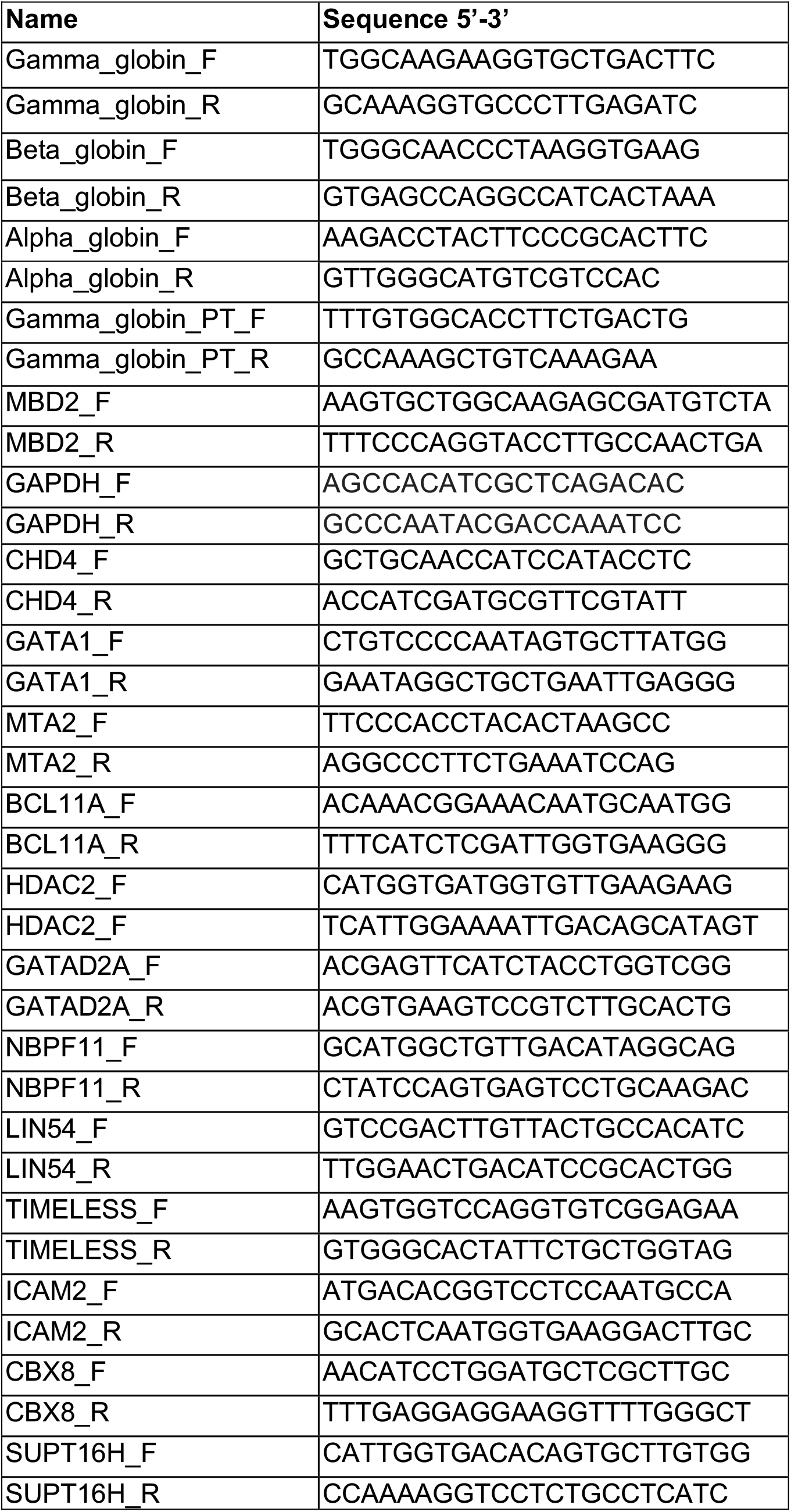
Primers for RT-qPCR

**Table S2.**
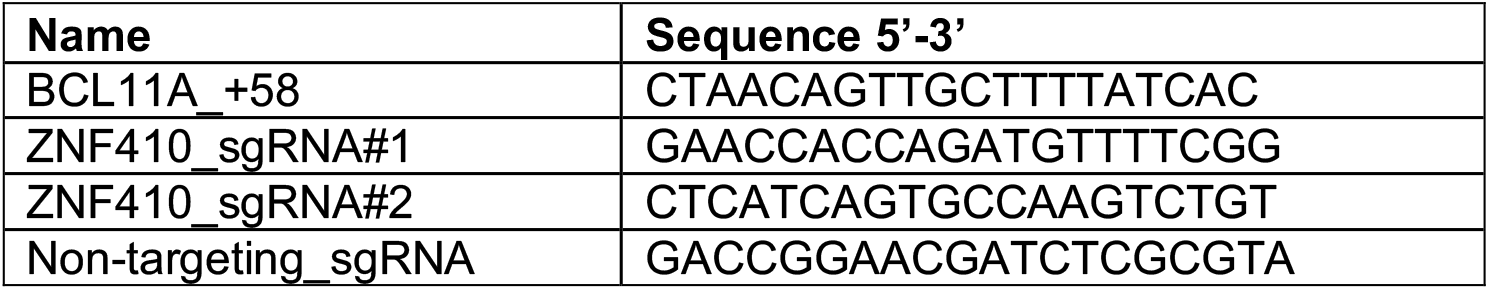
sgRNA sequences

**Table S3.** Plasmids used in this study

| **Plasmids** | **Vector** | **Tags** |
| --- | --- | --- |
| pSDM-HA-ZNF410 | pSDM101 | N-HA |
| pSDM-HA-CHD4 | pSDM101 | N-HA |
| pSDM-HA-ZF(1-5) | pSDM101 | N-HA |
| pSDM-HA-ZF(2-4) | pSDM101 | N-HA |
| pcDNA3-FLAG ZNF410 full-length 1-478 aa | pcDNA3 | N-FLAG |
| pcDNA3-FLAG ZNF410 ZF (1-5) 209-366 aa | pcDNA3 | N-FLAG |
| pcDNA3-FLAG ZNF410 ZF (1-4) 209-338 aa | pcDNA3 | N-FLAG |
| pcDNA3-FLAG ZNF410 ZF (2-5) 244-366 aa | pcDNA3 | N-FLAG |
| pcDNA3-FLAG ZNF410 ZF (1-3) 209-308 aa | pcDNA3 | N-FLAG |
| pcDNA3-FLAG ZNF410 ZF (2-4) 244-338 aa | pcDNA3 | N-FLAG |
| pcDNA3-FLAG ZNF410 ZF (3-5) 274-366 aa | pcDNA3 | N-FLAG |
| pGEX-6P-ZF (1-5) | pGEX-6P-1 | N-GST |

**Table S4.** Antibodies used in flow cytometry

| **Antibody** | **Clone** | **Source** | **Catalog#** |
| --- | --- | --- | --- |
| PE Anti-Human CD49d | 9F10 | BioLegend | 304304 |
| PE/Cy7 Anti-Human CD71 | CY1G4 | BioLegend | 334112 |
| APC Anti-Human Band3 |  | New York Blood Center |  |
| Anti-Fetal Hemoglobin-APC, human | HBf-1 | Thermo Fisher Scientific | MHFH05 |
| V450 Mouse Anti-Human CD45 | HI30 | BD Biosciences | 560367 |
| FITC Anti-Human CD235a | HI264 | BioLegend | 349104 |
| PE Anti-Human CD33 | P67.6 | BioLegend | 366608 |
| APC Anti-Human CD19 | HIB19 | BioLegend | 302212 |

**Table S5.**
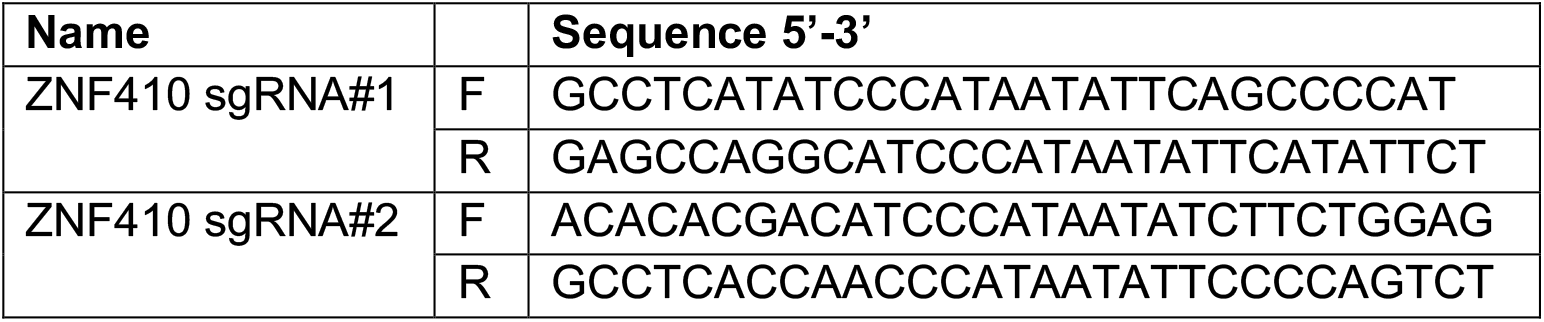
Primers for NGS sequencing

**Table S6.**
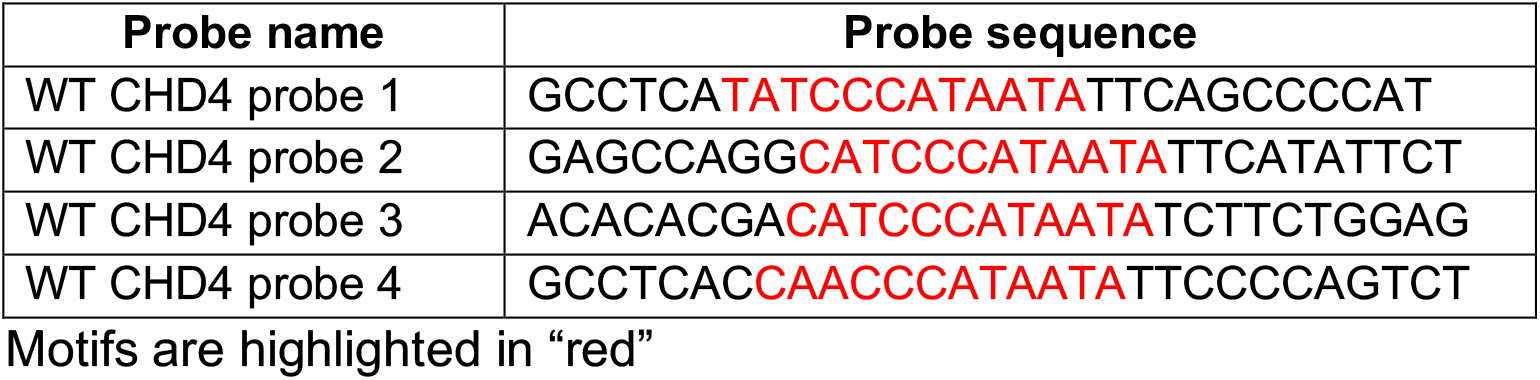
EMSA probe sequences

**Table S7.**
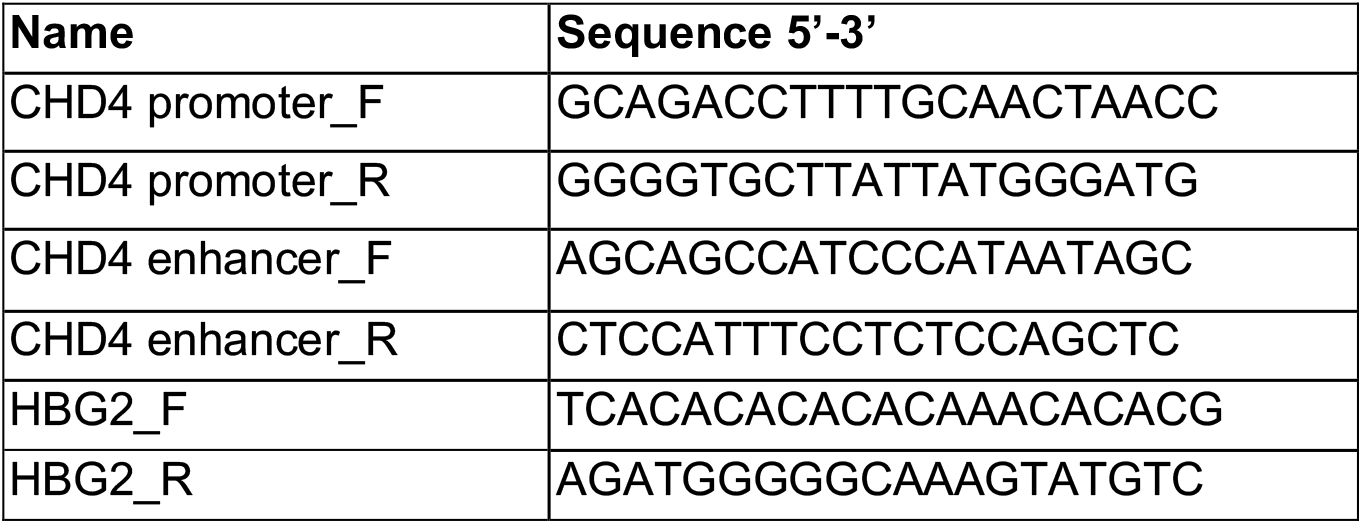
Primers for ChIP-qPCR

**Table S8.** Top20 differentially expressed transcripts from HUDEP-2 and primary CD34+ cells RNA-seq

| HUDEP-2 cells |  | Primary CD34+ cells |  |
| --- | --- | --- | --- |
| **Top10 upregulated genes** | log2(Z#2/NT) | **Top10 upregulated genes** | log2(Z#2/NT) |
| **HBG1/2** | 4.28 | HSPB1 | 2.15 |
| AC104389.5 | 4.01 | IFI27 | 1.78 |
| RNU5B-1 | 3.60 | **HBG1/2** | 1.75 |
| SNORD97 | 2.29 | SELENOW | 1.67 |
| CKB | 2.24 | RN7SL471P | 1.60 |
| PRG2 | 2.17 | PKN1 | 1.43 |
| H1F0 | 2.13 | AC104389.5 | 1.43 |
| AK1 | 1.83 | STUB1 | 1.40 |
| RNU5E-1 | 1.82 | PAFAH1B3 | 1.40 |
| CD63 | 1.75 | CDKN1A | 1.38 |
|  | log2(Z#1/NT) |  | log2(Z#1/NT) |
| **HBG1/2** | 4.26 | HSPB1 | 2.01 |
| AC104389.5 | 3.95 | SELENOW | 1.55 |
| CKB | 2.44 | **HBG1/2** | 1.47 |
| H1F0 | 2.41 | STUB1 | 1.46 |
| PRG2 | 2.09 | RPS19BP1 | 1.42 |
| AK1 | 1.99 | PAFAH1B3 | 1.35 |
| MIF | 1.92 | ISOC2 | 1.34 |
| CD63 | 1.90 | PKN1 | 1.32 |
| TMSB10 | 1.79 | COA4 | 1.30 |
| ATF5 | 1.65 | DYNLT1 | 1.27 |
| **Top10 downregulated genes** | log2(Z#2/NT) | **Top10 downregulated genes** | log2(Z#2/NT) |
| FOS | -1.88 | AC130456.3 | -2.03 |
| S100A10 | -1.59 | CBX6 | -2.01 |
| **AC092490.1** | -1.50 | TMCC2 | -1.69 |
| SLC2A3P1 | -1.49 | **AC092490.1** | -1.47 |
| SCARNA3 | -1.30 | ACSL6 | -1.47 |
| **CHD4** | -1.23 | KDM7A | -1.32 |
| SCARNA18 | -1.20 | ZNF410 | -1.24 |
| RNU6-11P | -1.11 | SMOX | -1.24 |
| TUBA1A | -1.07 | PRDX5 | -1.09 |
| PLA2G6 | -1.01 | **CHD4** | -1.08 |
|  | log2(Z#1/NT) |  | log2(Z#1/NT) |
| RF00019 | -2.43 | TMCC2 | -2.01 |
| S100A10 | -1.60 | ACSL6 | -1.67 |
| **AC092490.1** | -1.45 | CBX6 | -1.56 |
| FOS | -1.32 | **AC092490.1** | -1.50 |
| TMCC2 | -1.18 | ZNF410 | -1.49 |
| GLIPR2 | -1.14 | SMOX | -1.45 |
| TUBA1A | -1.06 | PRDX5 | -1.00 |
| **CHD4** | -0.99 | **CHD4** | -0.93 |
| PPP4R1 | -0.95 | SPC25 | -0.93 |
| SCARNA3 | -0.89 | GPCPD1 | -0.93 |

**Table S9.**
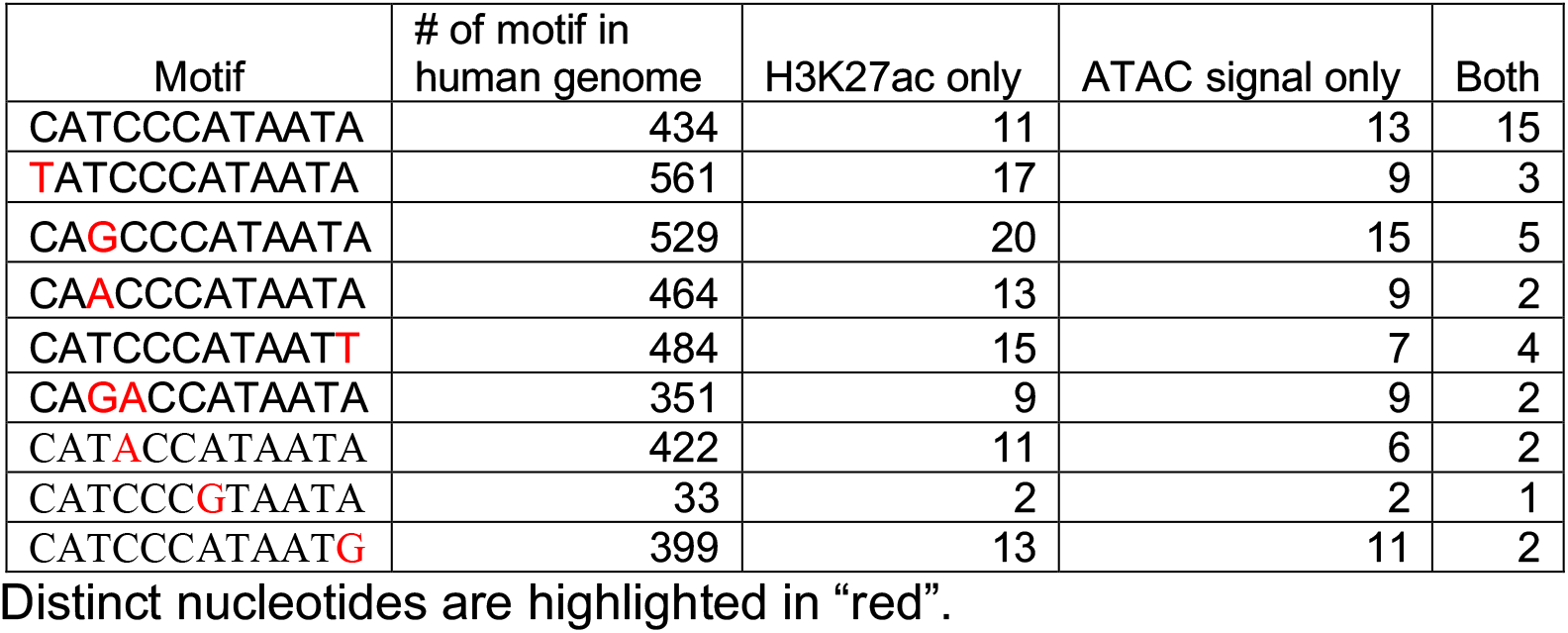
Analysis of H3K27ac ChIP-seq and ATAC signal from primary human erythroid cells at ZNF410 motifs

| Motif | # of motif in human genome | H3K27ac only | ATAC signal only | Both |
| --- | --- | --- | --- | --- |
| CATCCCATAATA | 434 | 11 | 13 | 15 |
| **T**ATCCCATAATA | 561 | 17 | 9 | 3 |
| CA**G**CCCATAATA | 529 | 20 | 15 | 5 |
| CA**A**CCCATAATA | 464 | 13 | 9 | 2 |
| CATCCCATAAT**T** | 484 | 15 | 7 | 4 |
| CA**G**ACCATAATA | 351 | 9 | 9 | 2 |
| CAT**A**CCATAATA | 422 | 11 | 6 | 2 |
| CATCCC**G**TAATA | 33 | 2 | 2 | 1 |
| CATCCCATAAT**G** | 399 | 13 | 11 | 2 |

Distinct nucleotides are highlighted in “red”.

**Table S10.**
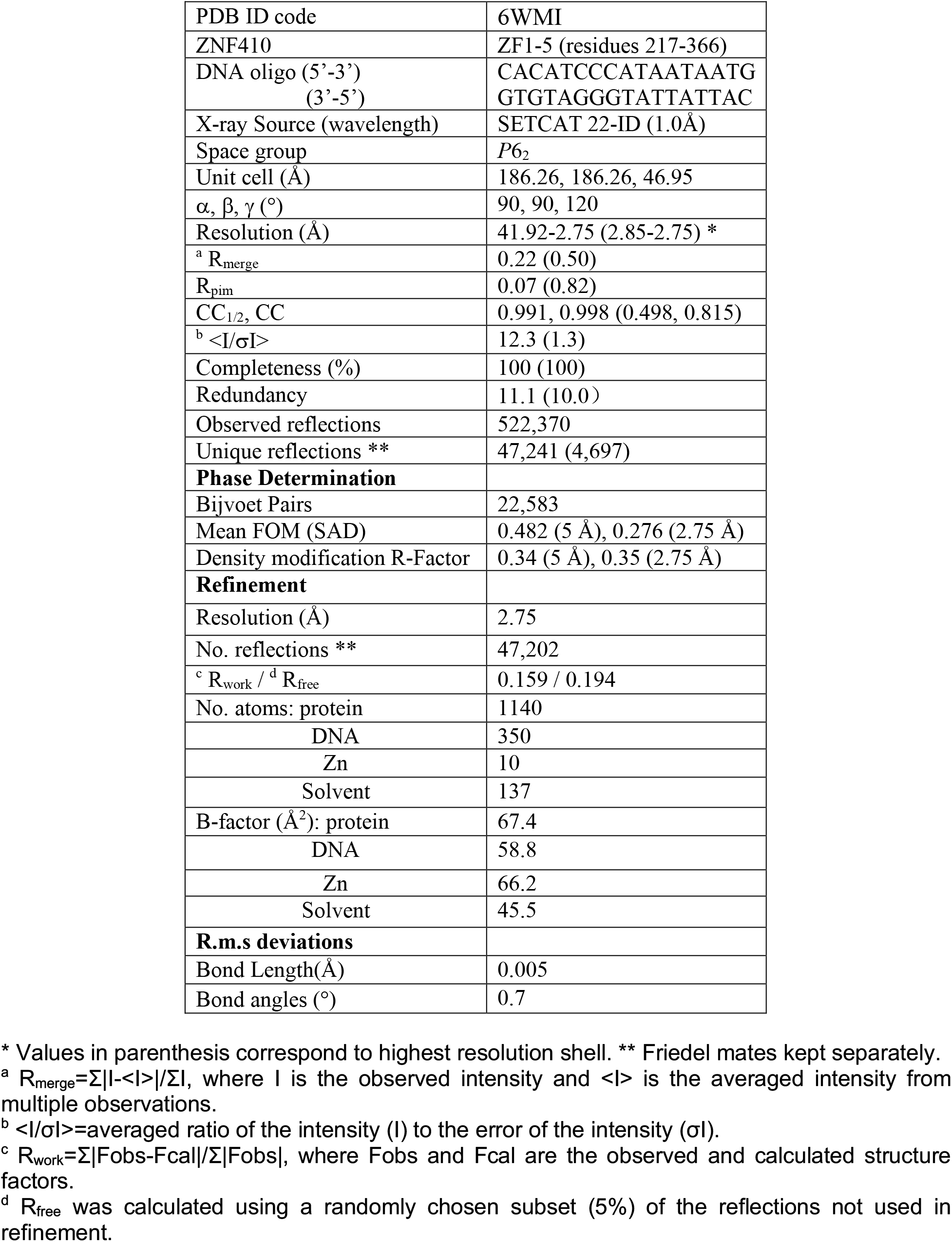
Summary of X-ray data collection and refinement statistics

